# Media matters: A resource for understanding the determinants of migratory potential of malignant and non-malignant cells

**DOI:** 10.64898/2026.11.18.767665

**Authors:** Bareun Kim, Anthony T. Lopez, Maria F. Osuna, Indhujah Thevarajan, Sonia Fuentes Lewey, Eustolia Sauceda, Dorothy Mundy, Chendong Yang, Lei Wang, Nigel Cole, Keala Minna-Choe, Isaiah Sanchez, Urooba Nadeem, Boning Gao, John D. Minna, Mary Wight-Carter, Elma Zaganjor, Ralph J. DeBerardinis, Jihan K. Osborne

## Abstract

Cell mobility is critical for homeostatic processes such as wound repair and diseases like cancer metastasis. While the advent of migrastatic and antimetastatic therapies has promise, the identification and translation of pathways important for migratory plasticity into druggable targets approved in clinical trials are considerably lacking. To better understand why, we developed an unbiased high-throughput wound-healing/invasion assay to screen for the migratory potential of >60 non-malignant and malignant cell lines derived from the pancreas, colon, skin, breast, prostate, and lung. The goal of this phenotypic screen was to provide a resource of migration/invasion potential and insight into the differential properties not only between malignant and non-malignant cells but also the various carcinomas. Surprisingly, we found non-malignant epithelial cells of the lung, skin, and pancreas migrated/invaded faster than their malignant counterparts. We observed perturbation of glucose metabolism, via inhibition of glycolysis, restricted the migration of non-malignant but not cancer cells, whereas inhibition of oxidative phosphorylation limited the mobility of malignant more than normal cells of the lung, skin, and pancreas. Short-term acclimation of cancer cells in low-nutrient media, containing low/no serum, low calcium, and physiological concentrations of glucose, increased in vitro migration/invasion and in vivo metastatic potential of lung but not pancreatic carcinomas. Phenotypic and metabolic screens presented here could identify prospective therapeutics not only for cancer metastasis, but also diseases driven by the dysregulation of wound-healing/regeneration.

**One Sentence Summary:** While phenotypic screening indicated normal cells migrated faster than their cancer counterparts; lung cancer cells exhibited greater metabolic plasticity to low glucose conditions, enhancing migration and invasion in vitro and metastasis in vivo.

## Introduction

Different modes of cell migration noted to occur in vitro, on 2-dimentional and/or in 3-dimentional (2-D, 3-D) substrates are linked to cellular properties that when dysregulated, correlate with increased pathology in vivo. These descriptive modes incorporate parameters of cell function and shape/morphology regulated by mechanosensory, bioenergetics, and biochemical signaling that lead to gene expression and ultimately cytoskeletal rearrangements^1–4^. Cell migration, critical for morphogenesis^5–7^, wound healing/regeneration^8–14^ and metastasis, is proposed to require an augmented abundance of energy^12,15–20^. Given that recent studies demonstrate that shifting cell morphology not in vitro cell migration/invasion assays, are good predictors of metastatic potential, further investigations into factors that accurately translate in vitro studies in vivo are needed.

In vivo studies in the mouse, as well as numerous human pathology reports, have described diverse modes of migration used by epithelial cancer cells (i.e., carcinomas), ranging from the movement of a single cell (via mesenchymal and amoeboid modes) to the migration of structured collective epithelial sheets^7,21–24^. While induction of the metastatic cascade is often portrayed as the dissemination of individual cells from the primary tumor as a result of post-epithelial to mesenchymal plasticity (EMP) induction, cluster formation of cancer cells characterized by increased apical junctional complexes (E-Cadherin+ and Claudin+) has been demonstrated to invariably increase metastatic potential^2,25–28^. Similarly, normal non-malignant epithelial cells also display collective and single-cell modes of migration as their means of movement during embryonic development, wound healing and asthmatic bronchoconstriction^3,6,8,13,25,29,30^. Given these previous in vivo accounts, we designed a phenotypic screen that encompasses both single and various modes of collective migration, via high-throughput wound-healing on and in substrates such as collagen or Matrigel.

Our initial hypothesis for this mobility screen was founded on the premise from decades of literature, that robust cell migration in cancer cells (quantified by speed, increased number of cells migrating, variability of cell morphology, persistence, and track length when compared to normal cells), could be an accurate measure of metastatic potential. We conducted the unbiased migratory screen of 60+ cell lines to identify cell lines with high migration and invasion potential for further examination. Contrary to the prevailing theory, nearly all the normal/non-malignant cells from the lung, pancreas, skin (keratinocyte and melanocytes), and colon invaded/migrated more robustly than cancer/malignant cell counterparts. Thus, our questions became: ‘what are the fundamental differences between how cancer cells and normal cells migrate/invade and how does the extracellular environment control migration output?’.

Several recently published studies have elegantly revealed the complexity of metabolic flexibility, diversity, and plasticity of the tumor microenvironment (TME) and within the tumor itself during metastasis^31–34^. We previously published one potential solution for why normal lungs cells moved faster was due to increased basal and post-wounding ATP levels^35^. We screened a subset of non-cancerous and cancerous cell lines from lung, pancreas and skin in the same media. Low-nutrient exposed lung cancer cells increased their migration to the same extent as the non-cancerous lung cells. We tested our observations of differential migratory capacity in vivo using xenograft models of growth (subcutaneous tumor formation) and metastasis/colonization (intravenous tail-vein and orthotopic instillation). Solidifying our in vitro observations, we detected lung cancer cells pre-exposed to low nutrients colonized various tissues while high nutrient treated cells mostly colonized the lung. The observations of lung in vitro and in vivo were not universal across all epithelial cancers. Pancreas and breast cancer cells were either more dependent on high nutrients for proliferation/survival (pancreas) or had no effect (breast) further indicating a unique opportunity for further investigation into more personalized metastatic therapeutics depending on cancer type. Collectively, our manuscript provides a comprehensive resource as well as a cautionary tale for how to design and interpret in vitro migration assays to better recapitulate in vivo phenomena.

### Phenotypic screening reveals that normal cells migrate/invade faster than cancer cells

Using the SX5 Incucyte live-cell imager, we developed an unbiased high-throughput screen to identify cell lines with high migration and invasion potential (Figure1A). We adapted all human malignant cell lines to RPMI1640 medium supplemented with 5% fetal bovine serum (FBS; R5) for 8-10 passages, while all human non-malignant cells, (apart from MCF10A, normal colonic and normal melanocyte cells) were grown for 6-8 passages in keratinocyte serum-free medium (KSFM). On collagen coating, we screened malignant cell lines isolated from primary and metastatic tumors of lung, colon, pancreas, breast, skin (melanoma and basal cell-like carcinoma) and prostate and their non-malignant counterparts for the ability to migrate and/or invade collagen/Matrigel post-wounding in their respective growth medium (R5 or KSFM; Extended Data Table S1). To achieve a quantifiable unbiased readout of migration potential, we measured **<u>T</u>**ime for wound **<u>D</u>**ensity to reach 80%, or 50%, or 40%. – (TD80, TD50, or TD40) using the Incucyte analysis software. We were surprised to find that the non-malignant cells of the lung, pancreas, and skin (keratinocytes and melanocytes) had significantly higher migratory potential than malignant cells from their respective tissues (Figure 1B-1E; Figure S1A), while the normal colon and prostate cells tested did not (Figure 1F and Figure S1B). In our previous study^35^, we observed that non-cancerous MCF10A breast cells, screened in their normal growth media, migrated significantly slower than the MDA-MB-231 breast tumor cells in their normal growth media, as previously reported^36,37^.

**Figure 1.**
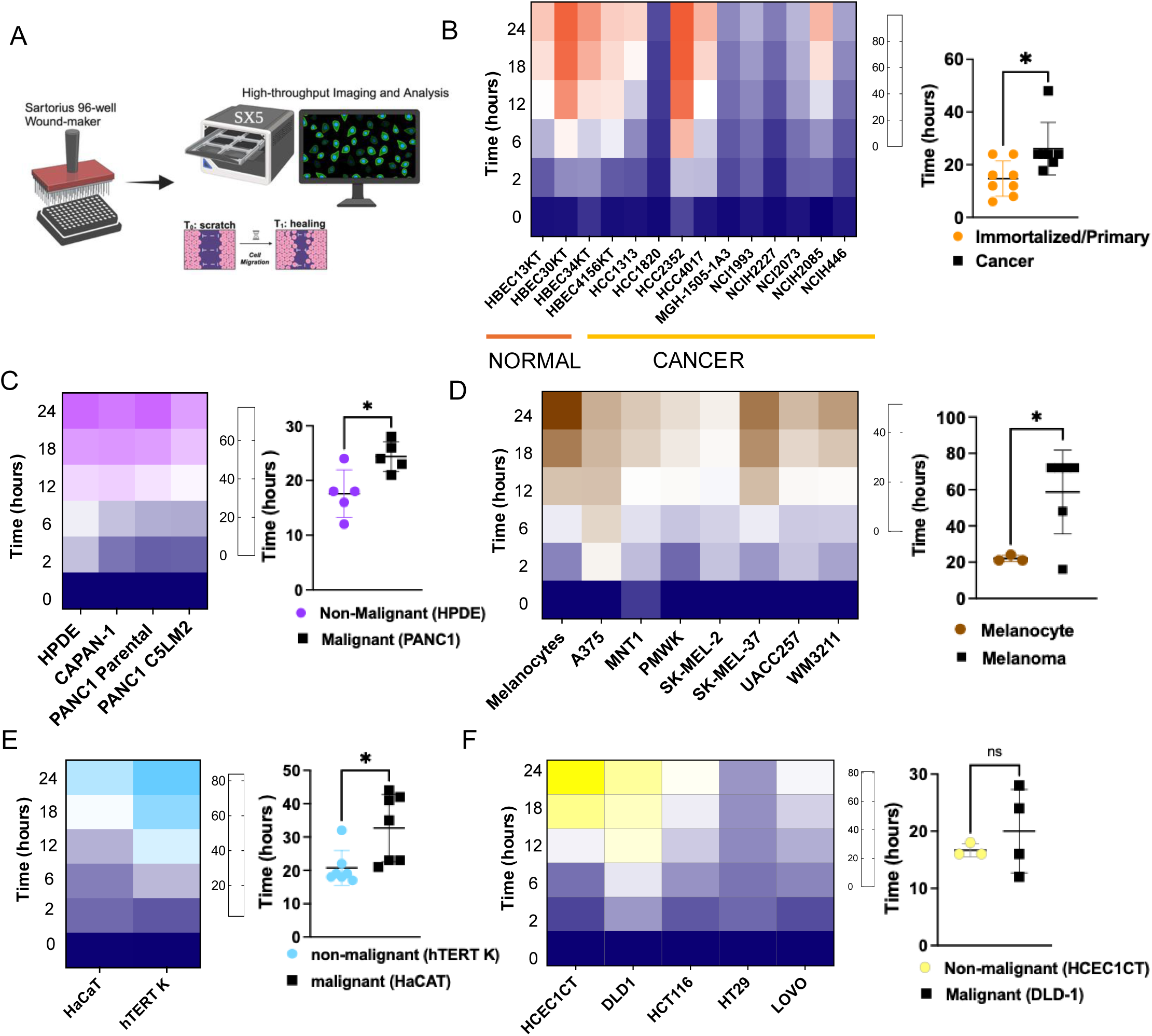
Non-malignant Cells Close the Wound Faster Than Malignant Cells. A) Schematic illustration of high-throughput migration/invasion screen using the SX5 Incucyte^®^ Live cell imaging instrument and analysis by the Sartorius cell migration software application. 50,000 cells were plated in collagen-coated 96-well plate (Corning, 3595) and/or Incucyte^®^ Imagelock 96-well Microplate (Sartorius, BA-04856). Wounds were induced in all wells using the Incucyte^®^ 96-well wound maker. B-F) Heatmaps (Left) depict the migratory rate of normal and cancer cells and dot plots (Right) represent the quantification of the time for wound of normal and cancer cells to reach at 80% B) lung (combined cell lines), C) pancreas, D) melanoma/melanocytes, E) keratinocytes and F) colon within 24-hour period. Statistical significance for B-F) are combined percentages where each dot represents single experiment of combined TD80 using Welch’s t-test. B) Graphs for normal lung cell lines are N=4 individual experiments for HBEC13KT, HBEC30KT, HBEC4156KT; N=3 for HBEC34KT; lung cancer cell lines are N=4 for HCC1313, HCC1820, HCC2352, HCC4017, NCIH1993, NCIH2073, NCIH2085; N=3 for H2227 and N=2 for MGH-1505-1A3 and NCIH446. *P=0.0208 C) Pancreas graph is from individual experiments for normal cell line, HPDE N=7 and PANC1 primary tumor cell line N=6. Heatmap data additionally represents PANC1-C5LM2 metastatic cell line N=8 and CAPAN-1 N=7. *P=0.021 D) Graphs for melanocytes N=3 and melanoma N=6. *P=0.011 E) Graphs of skin are from individual experiments normal cells, hTERT keratinocytes (N=7) and malignant cells, HaCaT (N=7). *P=0.021 F) Graphs for colon normal colon N=3 and N=4 DLD-1. ns=0.4316 All error bars represent standard deviations (SD). Panel A) made in BioRender.

Next, we selected several cell lines from the tissues above to further examine their ability to invade the extracellular matrix, Matrigel. Investigation into the ability of the lung cells to invade, confirmed the results we observed in migration assays (Figure S1C-S1D). Immortalized non-malignant bronchial epithelial cells were able to invade more than even the most invasive cancer cells tested in our screen (Figure S1C, S1D). However, immortalized normal pancreatic and pancreatic cancer cells both invaded to similar levels, while mammary and breast cancer cells differentially invaded as observed during 2-D migration (Figure S1E, S1F).

### Intracellular ATP localizations but not levels correlate with increased migration

Cell migration necessitates a considerable expenditure of energy in the form of intracellular ATP ^20,38^. We imagined one major distinction between how malignant and non-malignant cells closed wounds could be the bioenergetics of ATP. Recent studies show during anchor cell invasion in *C. Elegans,* protrusive forces generated via invadopodium F-actin polymerization fueled by intracellular ATP (iATP), is generated by glycolytic enzymes clustering with mitochondria at the cell’s leading edge^16,17^. We derived cell lines stably expressing a lentiviral FRET-based ATP sensor in several non-malignant and malignant cells (Figure S2A). This gave us the ability to examine spatiotemporal distribution of the iATP at wound induction within the first few hours of migration into the scratch. Our recently published data detailed a greater upsurge in iATP at wound induction in normal mammary and normal lung epithelial cells than their tumor counterparts^35^. At wound induction, (T=0) in the non-malignant immortalized lung (HBEC30KT) and pancreatic (HPDE) cell lines, we observed an increase in iATP. However, in the malignant lung (HCC4017) and skin (HaCAT) cell lines we also observed an increase in iATP, indicating levels alone could not account for augmented migration (Figure 2A). The distribution of iATP in the non-cancerous cells were evenly disseminated throughout the cell body (Figure 2B and Figure S2B). Intriguingly however, in the cancer cells, we detected the presence of dynamic clusters of iATP localized at lamellipodia-like structures at the leading edge as cells migrated into the wound (denoted by arrows at various time points) (Figure 2C, 2D and S2C). This suggests, while similar biochemical mechanisms are engaged during migration in normal and cancer cells, the bioenergetics of iATP localization may be a key measurable factor of robustness. However, we pondered that this would need to be further examined with malignant and non-malignant cells in the same media. We next reexamined several of our robust malignant and non-malignant migrators under the same culture conditions.

**Figure 2.**
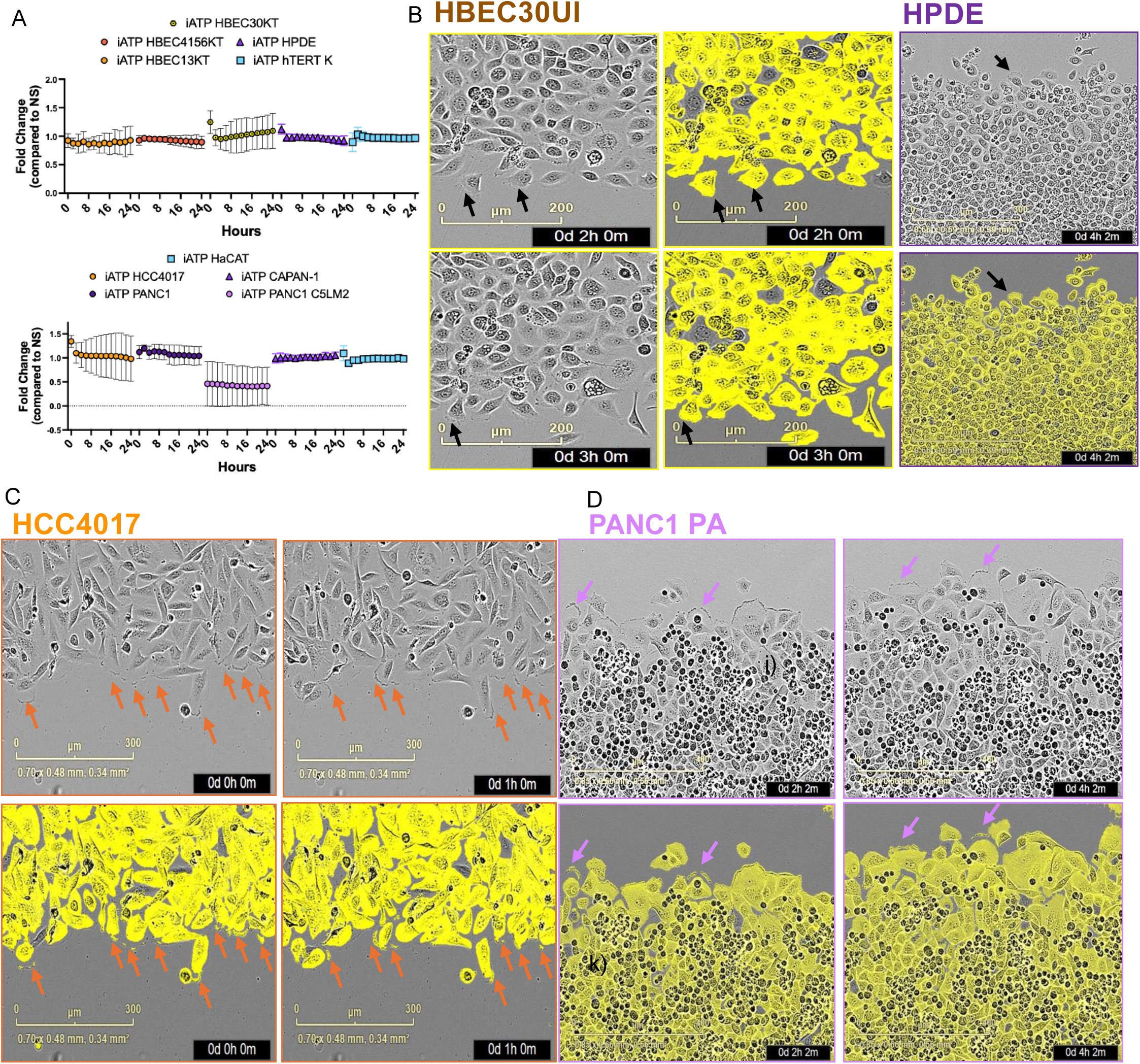
ATP is differentially localized in cancer and non-cancerous cells. Cells stably expressing lentiviral CytoATP vector were plated in 96-well collagen-coated Corning and/or Incucyte^®^ migration plates to confluence (50K). Wounds were induced in all wells using Incucyte^®^ 96-well wound maker then analyzed with the SX5 metabolism module and data analysis software. Time-lapse visualization of the cells, A) Fold change in iATP during migration were normalized to unwounded confluent cells. Graphs represent standard deviation of N=2 independent experiments. (B) HBEC30UI and HPDE C) HCC4017, and D) PANC1 PA during early time points of migration showing dynamic iATP-localized to the membrane ruffles of cancer but not normal cells. Arrows (black, violet and orange) mark iATP localization to the membrane ruffles of cancer cells. Representative images from duplicate wells of N=4 independent experiments.

### Low-nutrient conditions induce faster migration but slower proliferation

Before rescreening our robust migrators, we did initial testing on the serum concentration, conducting 3 different measurements 1) at early migratory events (2 hours), 2) at later migratory events to account for the time needed to reach TD80 (48 hours) and 3) the fold change (Fc) of the wound density from the initial migration event to the later migration event. We tested the HCC4017 NSCLC cell line as our initial model cell line for two reasons. First, HCC4017 is a matched malignant/non-malignant pair with the unimmortalized primary HBEC30UI, and the immortalized HBEC30KT (immortalized via CDK4 and hTERT overexpression) cell lines from the same 62-year-old smoker. This allowed us to test malignant, and non-malignant cells from an isogenic background. Second, we used HCC4017 because it was derived from a stage 1A lung tumor, meaning the tumor was localized to lungs and had not spread to lymph nodes or any other distant parts of the body^39^, so any colonization events we documented here would not be confounded by previous metastatic occurrences from the patient. We conducted short- and long-term acclimation of HCC4017 to the non-malignant cell media, KSFM (which we used for growth of the HBEC30 cells) and RPMI 1640 supplemented with 2%, 5%, and 10% serum. There was no significant difference in wound density when we compared serum concentrations in RPMI1640 at early or late time points (Figure S3A-C). Long-term treatments were ≥ to 2 weeks and short-term treatments lasted exactly three days. Of note the short-term R10 treatments induced a greater Fc in wound density than R5 (Figure S3D). Short-term R10 exposure when compared to long-term R10 treatments, showed significantly less of a Fc in wound density but were not statistically significantly from the Fc of the K2 treated cells (Figure S3D). Thus far, we concluded the short-term R10 treatments of the HCC4017 cells had the greater capacity for wound closure but only acutely, as long-term treatments decreased wound density when compared to all other concentrations of serum. While the acutely treated R10 cells also had greater capacity for wound closure, we could not rule out the proliferative advantage that 10% serum could have.

To rule out the contribution high serum could have on proliferation during migration, we investigated proliferation in the various serum contributions. We have previously reported the doubling time of HCC4017 grown in R5 to be close to 20 hours^35^. At the earlier time points between 24-48 hours, we found all concentrations of serum R2-R10, and K2 doubled similarly (Figure S3E). While K2 cells had a high rate of wound closure, this did not translate to an apparent growth advantage (Figure S3C and Figure S3E). Taken all together, we found serum concentrations within the same media have miniscule effects on short-term proliferation (less than 48 hours), while long term growth defects were observed in the R2 and K2 conditions (greater than 48 hours), this contrasted with our data on migration where both lower concentrations of serum were able migrate similarly (Figure S3C-D). Thus, we concluded short-term treatments in the lower nutrient conditions found in R2 and K2 conferred less of a growth/proliferative advantage but were suitable for comparisons of migration potential as there was not much difference in the cell’s ability to close the wound, regardless of duration of treatment or effect on proliferation (Figure S3A-E).

Finally, we conducted a factorial reduction experiment, in four cancer cell line models of lung (HCC4017), pancreas (PANC1 parental and the liver metastatic clone, PANC1-C5LM2) and skin (HaCAT) starting with RPMI1640 without glutamine or glucose (R0). Short-term acclimation in R0 resulted in cytostatic and migrastatic effects in HCC4017 and HaCAT, with no observable defects in mobility in either of the pancreatic cell lines (Figure S3F-3I). Addition of 5mM glucose (R5G)–the same concentration found in people with normal glycemic indices^40^– led to a nearly 4-fold change in migration of HCC4017 and a 5-fold change of HaCAT cells relative to R0 (Figure S3F-3G). Again, no change was detected in either pancreatic cell line treated with R5G (Extended Data Figure S3H-3I). Interestingly, addition of 11mM glucose (R11G)–a supraphysiological concentration, equivalent to a person with hyperglycemia–significantly decreased migration in the lung cancer cell line (Figure S3F) or had no effect on migration in any of the other cell lines (Figure S3G-3I). The addition of 5% serum to R0, R5G, or R11G did not further potentiate movement among the lung, skin or primary pancreatic cancer cell lines, however there were significant changes in migratory potential in metastatic pancreatic cells relative to the R0 cells with 5% serum (Figure 3I). PANC1-C5LM2 cells in R5 media without glucose or glutamine significantly increased their migration 2-fold and 3-fold with the addition of 5mM and 11mM glucose respectively (Figure S3I). A previous study indicated cells can couple the energy used for invasion and proliferation from the same ‘energy depot’ ^41^, they also further demonstrated there is a larger amount of energy invested in proliferation than migration. Thus, this could mean under low glucose conditions, it would be actually advantageous for the cell to invest energy in migration/invasion over proliferation^20,41^. We found our data to be in large part aligned with these previous studies, indicating while a certain quantity of glucose (within euglycemic range) is necessary to provide the ATP/energy needed for migration, too much hindered migration/invasion.

**Figure 3.**
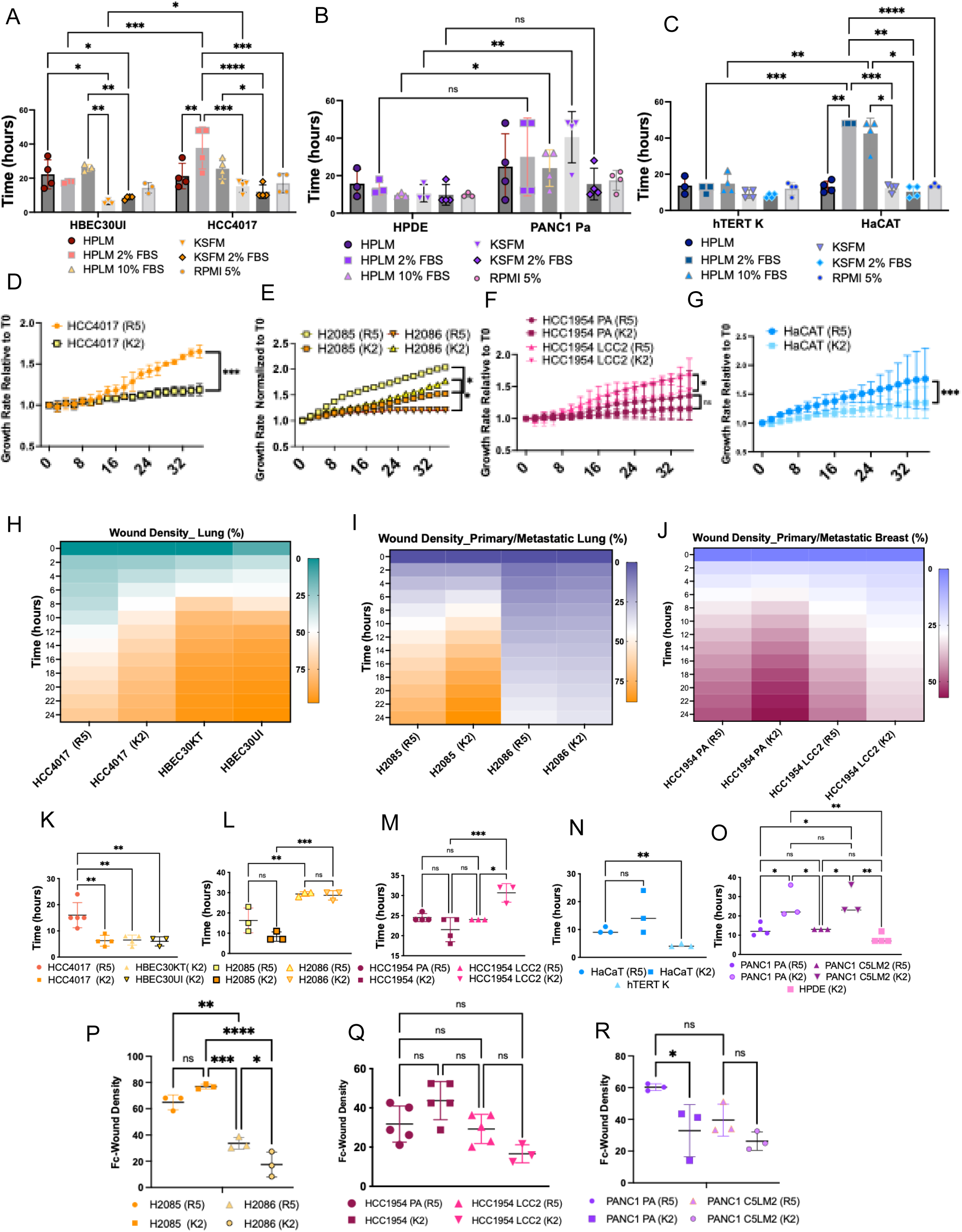
Low-nutrient Conditions Induce Faster Migration But Slower Proliferation. TD50 quantification of wound density of cells adapted short-term to human plasma-like media + 2% and 10% FBS (HPLM), KSFM +/− 2%FBS (K2) or maintained in RPMI 1640 5%FBS (R5). Statistical significance achieved by two-way ANOVA/mixed model analysis of individual experiments, A) HBEC30UI N=3/HCC4017N=4 (*<0.03, **<0.008, ***<0.005, ****<0.0001); B) HPDE N=3/PANC1 N=4 (*<0.03, **<0.005); C) hTERT K N=4/HaCAT N=4 (*<0.03,**<0.003, ***<0.0006). Graphs showing the growth rate of cells adapted short-term in K2 (low glucose/low serum/lower glutamine) or maintained in R5 (higher glucose/high serum/high glutamine) media relative to time zero of D) HCC4017 E) NCIH2085 and NCIH2086 F) HCC1954 PA and HCC1954 LCC2 G) HaCAT for 36 hours. Statistical significance achieved by two-way ANOVA of individual experiments, HCC4017 N=3 (***P=0.0002, 16-36 hours); HCC1954 N=3 (*P<0.0370, 12-16 hours, LCC2); HaCAT N=3 (**<0.006, 24-36 hours) H2085/H2086 N=2 (*P<0.0211, 10-36 hours); and ns=not significant (>0.83) Heatmaps showing the wound density of cells adapted to K2 or maintained in R5. H) HCC4017 and normal lung cell lines HBEC30KT and HBEC30UI, I) human primary lung cancer cell line NCIH2085 and metastatic lung cancer to lymph nodes line NCIH2086. J) human primary breast ductal carcinoma cell line HCC1954 PA and latent brain metastasis HCC1954 LCC2 24 hours after wound induction. K-O) Graphs measuring TD50 of non-malignant and malignant cell lines adapted to K2 or R5 K) HCC4017, HBEC30KT and HBEC30UI, L) NCIH2085 and NCIH2086, M) HCC1954 PA and HCC1954 LCC2, N) HaCAT and hTERT K O) Panc1 PA, PANC1 C5LM2 and HPDE. Graphs showing the quantification of fold-change (Fc) in wound density from 2-24 hours for P) NCIH2085/H2086 Q) HCC1954 PA/LCC2 R) PANC1 PA/C5LM2. Statistical significance achieved by one-way ANOVA of individual experiments N=3 for lung and pancreas and, N=4 for breast, *<0.05,**0.001, ***0.0001, ****<0.0001

We reassessed robust malignant and non-malignant migrators from lung, breast, pancreas, and skin in the same media and serum concentrations to compare their migratory potential in the same extracellular environment. We conducted short-term acclimation of all cells to the non-malignant cell growth medium–KSFM, the physiologic nutrient media^40^, human plasma-like medium (HPLM) or the cancer growth media–RPMI 1640 supplemented with 5% FBS (R5). Although, our initial screen of cancer cells was conducted in R5 and normal non-malignant cells in KSFM we acknowledge that different medias have vastly different components to support growth. Non-malignant cell growth media–KSFM–contains no serum (although supplemented with bovine pituitary extract), half the concentration of glucose (6.0mM) compared to the 11mM in RPMI1640 and interestingly have very similar concentrations of calcium (Figure S3J). Unfortunately, HPLM alone (which contains similar glucose, but 5x greater calcium than KSFM) supplemented with 2% or 10% FBS, did not support growth and/or migration of several of our malignant and non-malignant cell lines, apart from the skin cells hTERT K and HaCaT (only without serum) (Figure S3K). TD50 measurements of KSFM compared to HPLM and RPMI1640 revealed, KSFM supported the most robust migration, in the lung cells followed by K2 in the lung and pancreatic cells (Figure 3A-3B), while cells migrating in HPLM, provided the weakest migration phenotype across all cell lines (Figure 3A-C). Examination of all media conditions across the six cell lines revealed K2 and R5 supported robust migration and more importantly viability during migration. Thus, to maintain viability during migration we compared R5 as our high nutrient media and K2 as our low nutrient media.

Like the serum experiment, K2 media maintained constant, albeit significantly slower rates of proliferation in lung (Figure 3D-E), breast (Figure 3F), skin (HaCaT) (Figure 3G) with no significant change in pancreatic (PANC1 PA and PANC1-C5LM2) (Figure S3L) cancer cell lines grown under short-term conditions. In addition to the lung cancer cell line HCC4017, we found another primary lung cancer cell line, NCIH2085, but not its patient-matched lymph node metastasis, NCIH2086, slightly decreased the time required for cells reach wound density of at least 50% after being adapted to K2 (Figure 3H-K). Unlike lung, carcinomas from serval other tissues/organs did not adapt to low nutrient conditions by increasing their migration (Figure 3L-O). The breast cell line HCC1954, but not its latent brain metastases HCC1954 LCC2, adapted to K2 similarly to HCC4017 and H2085 in two out of the four experiments conducted (Figure 3L-M). We observed the non-malignant lung and to a lesser extent the skin cells, but not the pancreatic cells slowed their migration potential in R5 media (Figure 3A-C). Additionally, we noted HCC4017 K2 treated cells not only migrated as fast as the isogeneic non-malignant immortalized and primary cell lines, HBEC30KT and HBEC30UI, but also changed morphology (Figure S3M). HaCAT and PANC1 cell lines slowed their migration into the wound with K2-adaptation (Figure 3N-O).

One additional observation we made was the difference between primary tumors of lung, pancreas, and breast (NCIH2085, PANC1 PA, and HCC1954) and their isogenic metastatic counterparts (NCIH2086, PANC1–C5LM2, and HCC1954 LCC2). A recent study conducted a large-scale analysis of the in vivo metastatic potential of 500 cancer cell lines, spanning over 20 different tumor types^42^. Data exploration of the cell lines found in their MetMap125 study revealed cells from distant sites, retained higher metastatic potential than those from primary tumor sites (Figure S4A). However, this study like several others also indicate that primary tumors carry more of the molecular/gene expression signatures linked to cancer hallmarks such as ‘invasion or poor clinical outcome’ ^42–45^, additionally we observed that many cell lines available were taken from metastatic, not primary sites (Figure S4A). We examined the Fc of the primary and metastatic cells in migrating in K2 or R5 to get an idea of how the wound density from the initial migration event to the later migration event changes with time. Primary lung tumor cells, H2085, retained increased migratory potential when directly compared to metastatic tumor cells from their isogeneic counterparts, H2086, when measured in the same media (Figure 3I and 3P). Primary pancreatic (PANC1) and breast cells (HCC1954) like the TD50 measurements showed no difference in their wound density rates when compared to metastatic counterparts (Figure 3Q-R and Figure S4B). Cumulatively these data indicate while lower nutrient conditions significantly slowed cancer cell proliferation in all cells tested (except for the pancreatic cells), migration was maintained or even in the cases of the lung, increased.

We next tested the invasion potential of the isogeneic cell lines HCC4017 and HBEC30UI/HBEC30KT. We found the non-malignant primary and immortalized cells HBEC30UI and HBEC30KT respectively both invaded Matrigel more than lung cancer cells tested in K2 and R5 (Figure S4C-D and MOVIE 1-3). Measuring the TD50 of invasion, both normal cell lines HBEC30UI/HBEC30KT, were faster than the cancer cells HCC4017, in R5 but not statistically different from the cells in K2 (Figure S4D). Taken together, these data suggest that high serum and non-physiological glucose concentrations slow the invasion of select malignant and non-malignant lung cells. Furthermore, normal lung cells were able to migrate and invade into the wound more than cancer cells, irrespective of media composition, as we previously reported^35^.

### iATP fluctuations in response to confluency and migration changes

Previous studies indicate during invasion/migration metastatic tumor cells enhance mitochondrial respiration as well as ATP production^32,46–48^. To investigate whether nutrient conditions change iATP, we again utilized our cell lines stably expressing the FRET-based ATP sensor. The levels of iATP at 100% confluency did not change when adapted to the various medias, however we observed non-malignant cells had significantly increased iATP when compared to malignant cells (Figure 4A). First, we examined whether iATP would change depending on the confluency of cells (Figure 4B-E). Sub-confluent lung cancer cells grown in K2 or KSFM, rapidly increased their iATP as the cells proliferated, while cells grown in R5 slowly increased their iATP (Figure 4B). Interestingly, low-confluent PANC1 cells did not change their iATP, regardless of the growth media used (Figure 4B). Moderate to highly confluent cancer cells decreased their iATP as cells decreased proliferation regardless of the media used (Figure 4D). Sub-confluent HBEC30 and HBEC14UI primary cells did not significantly change their iATP levels as they proliferated (Figure 4C), however like the cancer cells, moderately confluent HBEC cells also dramatically decreased their iATP as proliferation slowed (Figure 4E). Interestingly, both HBEC30 and HBEC14 primary unimmortalized cells grown in R5 cells slowed proliferation as previously described for other non-malignant HBEC cell lines^49^. Taken together, we observed both cancer and normal cells dramatically decreased their iATP as confluency increased. However, while sub-confluent lung cancer cells grown in low nutrient media, rapidly increased their iATP, pancreatic cancer cells did not. This suggests cancer cells, like normal cells do maintain the ability to sense neighboring cells via mechanisms that include iATP regulation. There were significant differences in how lung cancer cells responded to low-confluency when compared to pancreatic cancer cells irrespective of the growth media conditions, which most likely reflect differences in physiology of the tissues of origin.

**Figure 4.**
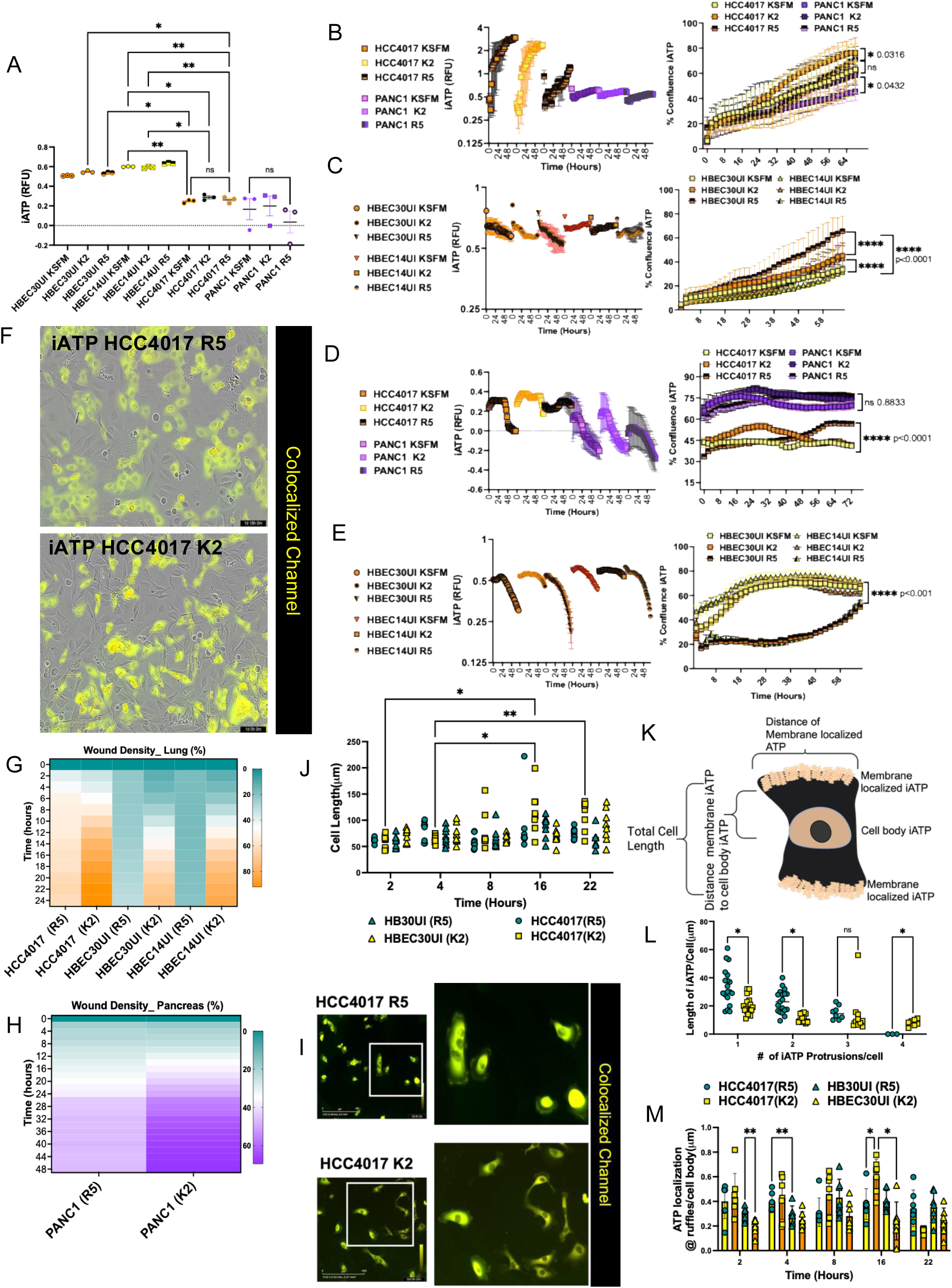
Quantification of proliferation and iATP localization in response nutrient changes. A) Measurement of iATP in normal (HBEC30UI, HBEC14UI) and cancer cells (HCC4017,PANC1) at confluence in various medias. Primary non-malignant (HBEC30UI and HBEC14UI) and malignant (PANC1 and HCC4017) stably expressing CytoATP – iATP construct were plated to 100% confluently. iATP levels were measured. Statistical significance for A) was achieved by one-way ANOVA of, Each dot represents individual experiments, N=3 individual experiments. B) Malignant (PANC1 and HCC4017) and C) primary non-malignant (HBEC30UI and HBEC14UI) cells stably expressing CytoATP construct were plated sub-confluently (<20%) and more confluently cells D) Malignant cells-PANC1 (>60%, <75%) and HCC4017 (>30%, <45%), E) HBEC30UI and HBEC14UI (>20%< 40%). Changes in iATP were measured in all cell lines B-E), for 48hours during adaptation to medias – KSFM, KSFM+2%FBS (K2), and RPMI 5%FBS (R5) while proliferation was simultaneously measure in cell adapted for up 72 hours in KSFM, KSFM+2%FBS, and RPMI 5%FBS. F) Examination of gross morphology of HCC4017 cells expressing CytoATP – iATP sensor in R5 and K2. Heatmaps quantifying the % wound density post induction in the stably iATP expressing malignant and non-malignant G) lung cells H) pancreas cancer cells adapted to R5 or K2 over the course of 48hrs. I) Images of cyto-ATP expressing HCC4017 cells plated sub-confluent to images membrane dynamics of the iATP construct in R5 and K2. Differential localization of iATP HCC4017 grown in R5 or K2 media. J) Quantification of total cell length (μm). K) schematic of iATP localization for (J-M). Panel K) made in BioRender L) X-axis quantifies the number of iATP protrusions per cell, Y-axis quantifies length of iATP protrusions at the membrane (μm). M) Quantifies the dynamics of the distance of iATP between ruffles/protrusions at the membrane and the cell body (μm) throughout migration.

While we observed no variances in iATP quantities there were obvious differences in the morphology of the HCC4017 cells grown in K2 versus R5, thus we hypothesized this shape modification could affect the migration potential (Figure 4F). Differences in cell morphology are intricately linked to migration potential^1,50^. Thus, we tested whether the iATP sensor would interfere with the migration capacity under different nutrient conditions. Like the cells without the sensor, both non-malignant and malignant cells grown in low nutrients closed the wound faster than those tested in higher nutrients (Figure 4G). The PANC1 cells stably expressing iATP interestingly, also increased their migration in the K2 media, unlike the unmodified cells (Figure 4H). We next investigated how iATP dynamics change during migration. We found that none of the cells tested significantly altered their iATP levels throughout migration, although K2 iATP levels seemed to increase slightly (Figure S5A-B). Taken together these data show that there were no observable differences in iATP levels of cells migrating in high and low nutrients. Again, this data is resonant with seminal studies that state if the prerequisite ATP quantities necessary for migration are achieved the amount of ATP used can be independent of migration potential^20^. This implies that while cell migration necessitates a considerable expenditure of energy in the form of ATP, like proliferation when, where, and why that increase in iATP is achieved may not be limited to solely to nutrient availability^51^, indicating a greater need for more examination.

We observed during migration malignant cells, unlike non-malignant cells, exhibited dynamic clusters of iATP at lamellipodia-like structures at the leading edge as cells migrated into the wound (Figure 2C-D and S2C). Thus, we investigated how these clusters changed during migration in the different types of media. First, we compared the primary unimmortalized, HBEC30 and the HCC4017 cancer cell lines both in R5 and K2 (Figure 4I-M and S5C-D). HBEC30 and HCC4017 cells grown sub-confluently in K2 had shorter and less cell protrusions labeled by the iATP construct (Figure 4I and S5C, white arrow), while cells grown in R5 had pronounced iATP labeled lamellipodia-like structures (Figure 4I and S5D, red arrows). We measured total cell length (Figure 4J), the length of the iATP lamellipodia-like structures (Figure 4L) and lastly the distance from the iATP at the cell ruffles to the iATP at the cell body (Figure 4M), visualized in (Figure 4K). HCC4017 cells grown in K2 significantly elongated their total cell length during migratory events, compared to HCC4017 grown in R5 and HBEC30 cells (Figure 4J). Furthermore, R5 grown HCC4017 cells had significantly greater lengths of the iATP labeled lamellipodia-like structures than the K2 cells as well as more iATP labeled protrusions (Figure 4L). We found that the distribution of iATP in the non-malignant cells were evenly disseminated throughout the cell body (Figure 2B). A recent study found that glycolytic enzymes reside on waves of actin during migration, which localizes glycolysis to the cell membrane^52^. Finally, we examined the distance of iATP at the ruffles compared to the iATP at the cell body, we observed fluctuations in the distance of the iATP at the ruffles from the cell body as cells migrated irrespective of media used (Figure 4M). Cumulatively, these data indicate cells grown in R5 notwithstanding disease states had increased iATP labeled lamellipodia-like structures and shorter total cell lengths that could be manipulated via exposure to decreased nutrients to increased migratory potential. Previous investigations cited here suggests that bulk migration is uniquely fueled by iATP localized at plasma membrane protrusions as a result of the combined efforts of both glycolytic enzymes in conjunction with mitochondrial output ^16,17,52,53^.

### Metabolite analysis indicates metabolic deviations during cell migration

Given recent data from several labs across multiple human tumors demonstrating how glucose deprivation shapes the metastatic potential^46,54–57^, we examined how glucose metabolism varies during cell migration of malignant and non-malignant cells. To better understand our findings that the migration potential of certain lung cells could be reprogrammed via decreased nutrient availability, we screened metabolite enrichment throughout migration. We used the non-malignant primary cell line HBEC30UI and the malignant cell line HCC4017 in high and low nutrients at wound induction (T=0) and during migration (T=4) normalized cells unperturbed at confluence (no scratch, NS). Hierarchical clustering interestingly suggested increased generation of various fatty acids and phospholipid species in non-malignant cells at T=0 and T=4 (Figure S6). Pathway enrichment analysis also detected augmented intermediates from fatty acid biosynthesis and suppressed intermediates from glucose and amino acid metabolism in the non-malignant cells (Figure 5A and S6). HCC4017 cells grown in R5 had decreased glycolytic intermediates and increased intermediates from the TCA cycle and amino acid metabolism (Figure S6). HCC4017 cells adapted to K2 showed increased glycolysis/gluconeogenesis intermediates, while all cells showed decreased intermediates of purine metabolism throughout migration. Targeted metabolite analysis revealed several of the same metabolites were transformed in the same direction at T=0 and T=4 in both the non-malignant and cancer cells grown in K2 although the cancer cells grouped together in the hierarchical clustering (Figure 5B and Supplemental Figure 6). Taken together, these data suggest distinct metabolic pathway utilization during the cell mobility events of normal wound-healing when compared to cancer cell migration.

**Figure 5.**
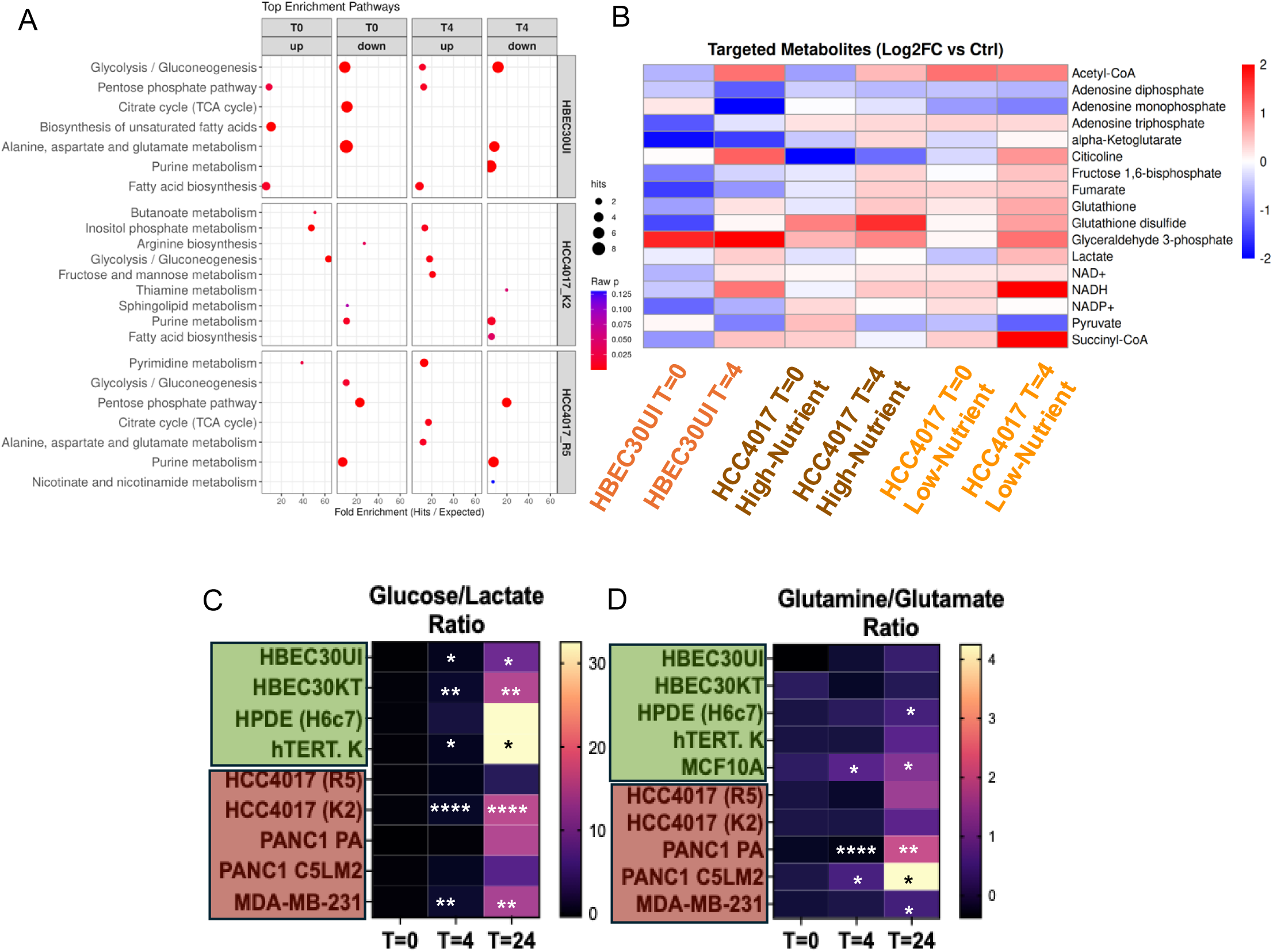
Metabolite analysis during migration. A) Dot blots of Top 3 enriched pathways from analysis of metabolomics data at wound induction (T=0) and during migration (T=4) normalized to control confluent plate of unscratched cells. B) Log-fold change of key metabolites identified from analysis at wound induction (T=0) and during migration (T=4) normalized to control confluent plate of unscratched cells. Heatmaps representing measurements using the NOVA analyzer of C) glucose consumption/lactate secretion and D) glutamine uptake/glutamine secretion into media at T=0, T=4, and T=24 normalized to NS. Non-malignant cells were tested in KSFM, while cancer cells in R5, except for the HCC4017 K2 cell line. Standard deviation of mean N=7 individual experiments for lung cells, N=3 pancreas, and N=3 breast cell lines. Statistical significance was assessed using a two-way ANOVA P*<0.05, **<0.005, ***<0.0005, ****<0.0001.

### Glucose metabolism diverges during migration in non-cancerous and cancerous cells

The tradeoff between glucose consumption and lactate secretion provides a means by which the ratios of their concentrations (Lactate/Glucose) can be used to estimate rates of glycolysis^58^. We detected a significant increase in glycolysis measured by increased lactate within the first 24 hours of wound induction from normal cells from the lung, skin, and pancreas but not cancer cells apart from MDA-MB-231 (Figure 5C). Interestingly, while we observed increased lactate production in our normal and K2-short term HCC4017 cells (Figure 5C) throughout migration; conversely, we simultaneously detected a significant increase in glutaminolysis measured by glutamine conversion to glutamate in the pancreatic and breast/mammary cells, but not the non-malignant lung and skin cells and the HCC4017 short-term adapted to lower nutrients (Figure 5D). Consequently, we measured the oxygen consumption rate (OCR), which primarily reflects mitochondrial respiration, and the extracellular acidification rate (ECAR), relevant to glycolysis. From our metabolite analysis, we observed decreased or no change in TCA/OxPhos intermediates in the non-malignant and K2-adapted cells (Figure 5B and Figure S6). We observed the basal rates of OCR were comparatively lower in the all the cancer cells after wound induction (Figure S7A). Like our previous data, the reverse was true of the basal rates of ECAR. After wounding there was no change apart from the R5-grown lung cancer cells which like the metabolomics data demonstrated lower glycolysis at wound induction (Figure S7B). In summary thus far, our data implies non-malignant and low-nutrient cancer cells produce lactate during migration, while high-nutrient cancer cells do not. This could indicate a shift towards glycolysis in our lung cancer cells adapted to low nutrient conditions similar to the non-cancerous cells grown under the same conditions.

Given the observation that K2-short term adapted HCC4017 cells significantly increased glycolytic rates compared to the R5 cells, we next examined whether suppression of glucose metabolism (via 2-deoxyglucose) or mitochondrial ATP synthesis (oligomycin) had any effect on cell migration. Before treating with the inhibitors, we evaluated expression of two of the five principal isoforms of hexokinase and the beta subunit of ATP synthase (ATP5B) in normal and cancer cells (Figure S7C-D). Hexokinase 1(HK1) is found to be ubiquitously expressed in almost all adult tissues while, hexokinase 2 (HK2) also is ubiquitously expressed, but at much lower levels than HK1; HK2 is notably expressed in highly glycolytic tissues, such embryonic and fetal cells and several cancers^59^. Expression of HK1 and low amounts of HK2 were detected in both normal and cancer cells throughout migration (Figure S7C-D), however, in the K2 media lung cancer cells increased the expression of both HK1 and HK2 (Figure S7C). Expression of ATP5B was low but still detectable in cancer cells four hours post-wounding during migration irrespective of nutrient levels (Figure S7C), while non-malignant primary cells had high levels of ATP5B throughout migration regardless of media (Figure S7D).

We determined the IC50 for migration potential of both 2DG and oligomycin in a range of normal and cancer cells from the same tissues. Normal cells from lung, pancreas, and skin had lower IC50 values than cancer cells in 2DG, with inhibition of migration occurring below 5mM in normal lung cells and 2.5mM in normal pancreas, and skin cell (Figure 6A, S7E and S7G). However, cancer cells in oligomycin had lower IC50 values than normal cells, with significant reduction in migration occurring below 2.5uM in malignant lung, pancreatic and skin cells (Figure 6B and S7F-G). Next, we examined whether the change in nutrient conditions could control the response to inhibition of glucose metabolism. Increased nutrients in the non-malignant and the malignant cells had slight but not significant effects on their response to 2DG or oligomycin (Figure 6C). Inhibition of migration in non-malignant cells in 2DG occurred between two-six hours post-wounding well below the observed doubling time whereas, reduced migration of malignant cells in oligomycin occurred between 24-28 hours closer to their observed doubling times (Figure 6C and S7I). Taken all together these data suggest non-malignant cells grown under low-nutrient/ physiologic conditions are sensitive to glycolytic inhibition during wound-healing; while migrating cancer cells in high nutrients are vulnerable to OxPhos suppression, while effects on proliferation of cancer cells cannot be completely ruled out.

**Figure 6.**
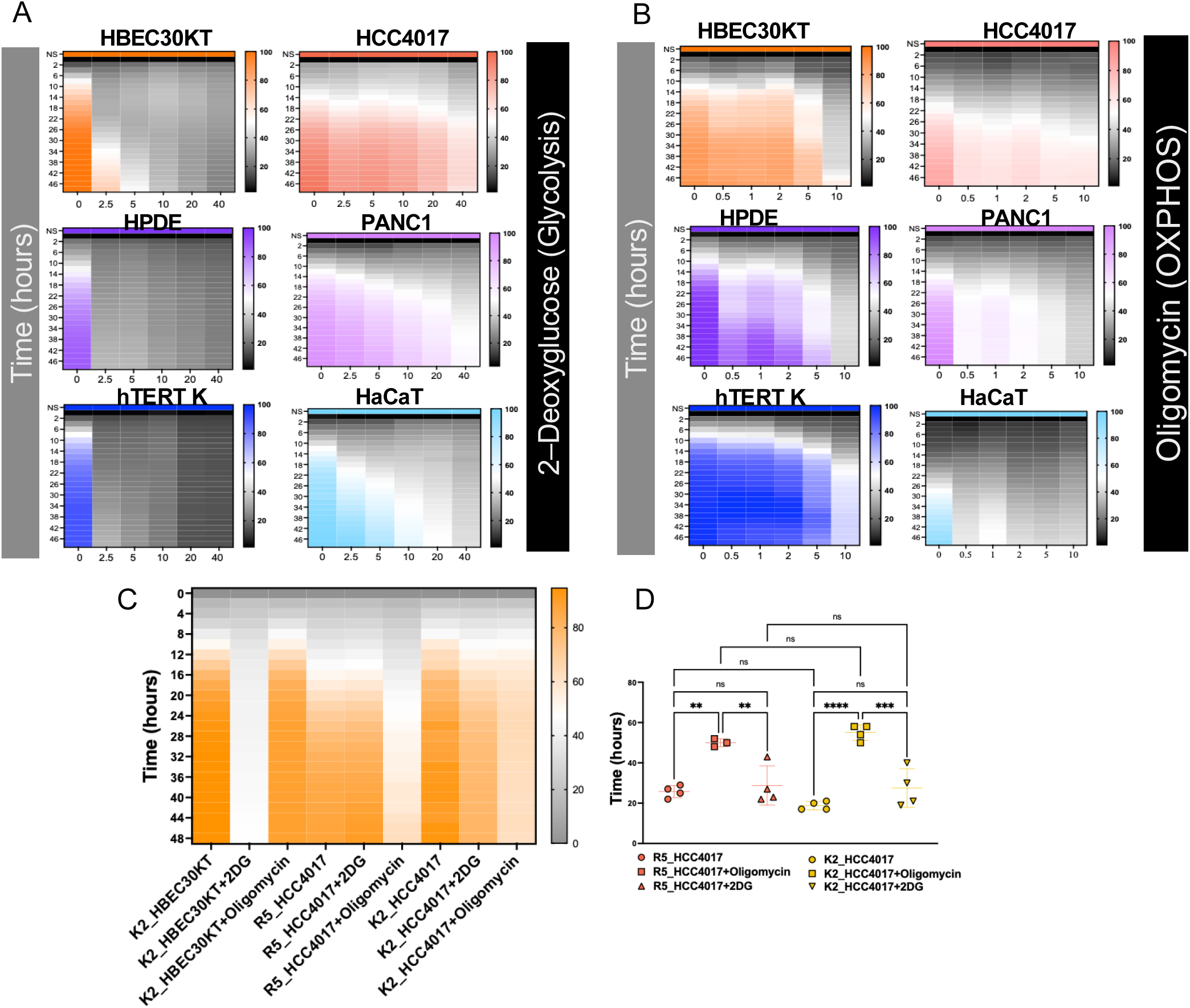
Divergent glucose metabolism during migration of non-cancerous and cancerous cells. Heatmaps quantifying the % wound density post induction. Quantification were made every 1-2 hours over the course of 48hrs of treatment with A) 2-Deoxy-D-glucose (2DG) or B) Oligomycin. Cell lines treated were from lung, pancreas, and skin. were treated with d) 2.5 mM, 5 mM, 10 mM, 20mM, 40mM, 2-DG and e) 0.5uM, 1uM, 2uM, 5uM, 10uM, Oligomycin. Non-malignant cells were tested in KSFM, while malignant cells in R5. HBEC30KT and HCC4017 cells were both adapted in K2 for 3-5-days or maintained in R5 media (HCC4017 alone). C)Heatmaps quantifying the % wound density post induction over the course of 48hrs of treatment with 0.5uM Oligomycin or 2.5mM 2DG. D) TD50 of panel C) R5 maintained or K2 adapted HCC4017 cells. Standard deviation of mean N=4 individual experiments. Statistical significance was assessed using one-way ANOVA P**<0.005, P***=0.0001, P****<0.0001.

### Lung cancer cells grown in low nutrients acquire more aggressive characteristics

Finally, to investigate whether the observations made in our in vitro experiments translated in vivo, we instilled R5 and K2 cells, subcutaneously (SubQ), intranasally (IN) and intravenous (IV) into Athymic nude mice (Figure 7). As previously stated, the HCC4017 cell line was originally isolated from a 62-year-old female smoker with a stage 1A lung tumor with no metastases present at diagnosis ^39^. This was an important criterion for our studies as even though the number of total primary tumors represented in the cancer cell line encyclopedia (CCLE) outnumbers the metastatic cell lines, when broken down there are more metastatic lung cancer cell lines derived from patients than primary lung tumors^42^. We introduced R5 and K2 cells from lung and pancreas, subcutaneously (SubQ) into the flank of Athymic Nude mice (Figure 7A and 7B). First, we developed SubQ tumors to assess tumor formation capacity. PANC1, grown in R5, were able to form tumors, while those adapted to K2 did not (Figure 7A). HCC4017 cells, regardless of nutrient conditions were able to form SubQ tumors to the same extent (Figure 7B), indicating short-term low nutrient exposure did not result in decreased capacity for tumor formation in vivo, as observed in pancreatic cancer cells.

**Figure 7.**
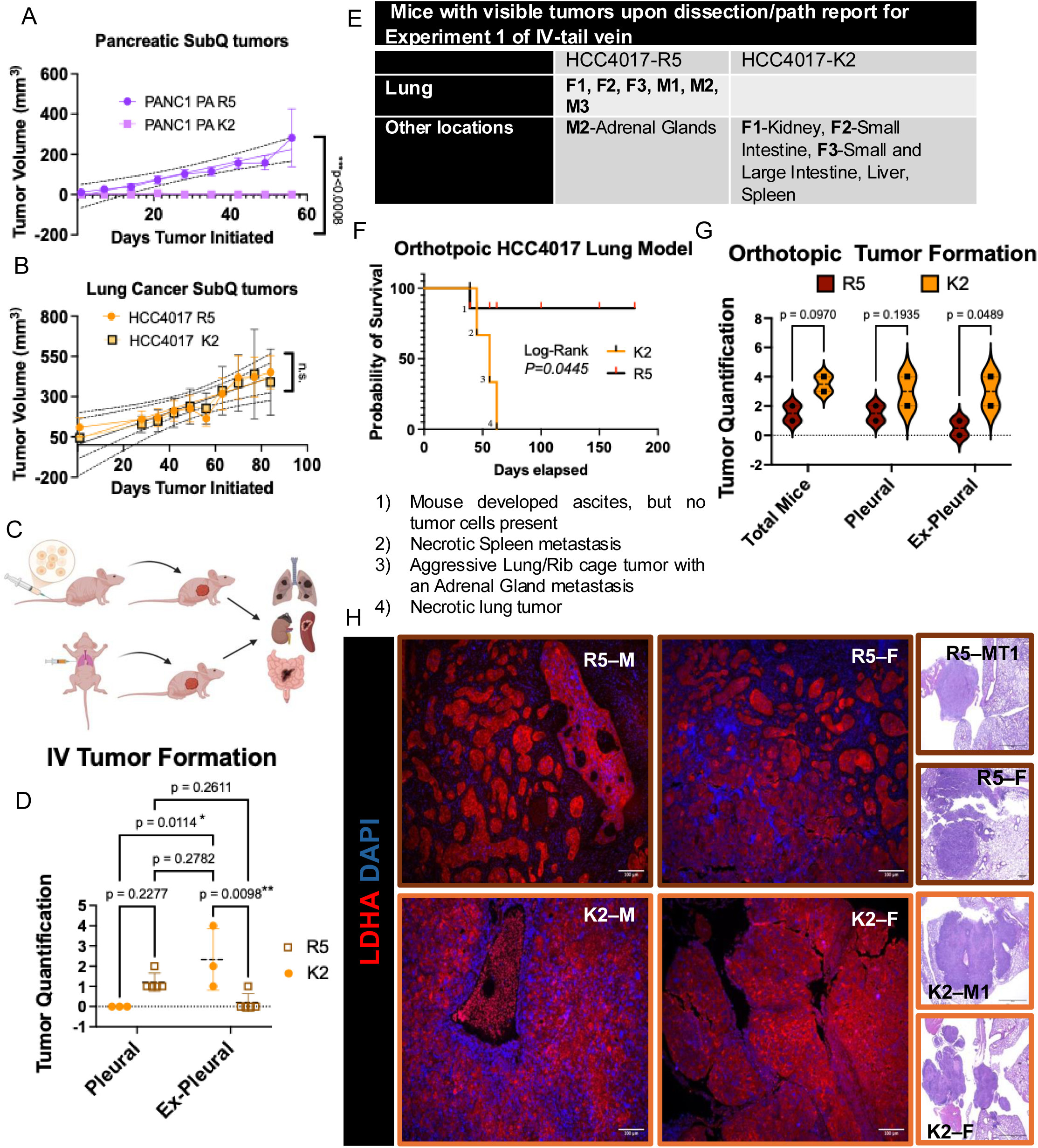
Highly glycolytic lung tumors have increased mortality rates in mice and human. Nude mice were injected with 1e6 A) HCC4017 cells or B) PANC1-PA in their normal growth media (R5) or adapted to low glucose/low serum (K2) for 3 days prior to being subcutaneously injected into flanks. N=10, 5 mice/group for HCC4017 and N=6, 3 mice/group for PANC1 PA. Tumor volumes were measured with electronic calipers. Means=SEM. P values for volume measurements calculated via linear regression (of slopes) in GraphPad. P for Lung, ns=0.8231, for Pancreas, ***0.0008. C) Schematic of two types of tumor cell administrations in Athymic nude mice used to measure metastasis, intravenously (IV) tail-vein top panel and orthotopic injections into lung bottom panel. D) Tumor formation of 2e6 HCC4017 cells administered by IV tail vein from adherent cultures. n=16 mice–R5, N=8; K2, N=8. Error bars represent SD of 2 independent experiments. Statistical significance assed via 2-way ANOVA. E) Table of tumor locations from experiment 1 of IV-tail vein. Oncospheres of cells derived from HCC4017 R5 previously formed tumors were either continuously grown in R5 media or adapted to K2 media for 3 days prior to orthotopic administration. Overall survival curves of mice bearing dissociated oncosphere adapted to R5 or K2 media. In K2 conditions, 3 out of 3 mice formed tumors and died as indicated on graph. In R5 conditions, 0 out of 3 mice died naturally, one mouse was sacrificed due to an ascites, however upon necropsy no tumor propagating cells were present in the liquid in the abdominal cavity. Presence of tumor cells assessed via organoid formation. The other mice bearing cells in R5 conditions were sacrificed 180 days post-surgical administration of cells. Statistical significance was assessed using log-rank test. P value indicated on panel. d) Quantification of two independent orthotopic lung models using Mixed-effects modeling (n=3 mice/ group) for first experiment and n=5 mice/group for second experiment. Multiple comparison tests were performed by controlling for FDR of q-values (p= 0.0970,p=0.1935, p=0.0481) of at least 0.05 by Two-stage set-up of Benjamini, Krieger and Yekutieli. e) Tumor-burdened lungs of a male and a female from K2-adapted and R5 from d) were formalin-fixed, sectioned and stained via immunohistochemistry (IHC) and IHC-immunofluorescence (IHC-IF) and for H&E and lactate dehydrogenase (LDH) and DAPI.

We then investigated the ability of cells to colonize/metastasize using two methods: 1) via IV tail-vein injections or intranasal instillation and 2) by orthotopic instillation directly into the left lobe of lung (Figure 7C). Prior to administering via IV, cells were infected transiently with an adenoviral luciferase construct to visualize short-term localization post-induction via bioluminescent imaging (BLI). Day 1 post-induction (PI) luciferase signals were mostly localized to tails and lungs, however by two weeks up to six weeks signal ceased in the body except in the tail (Figure S8A) Between 12-20 weeks PI, several but not all tumors grown in R5 and K2 became visible via BLI and X-Ray imaging (only in the lung) (Figure S8B-D). We found that cells grown in R5 developed lung tumors at a much higher rate than those that metastasized (Figure 7D-E and S8D). We observed cells acclimated to K2 formed lung tumors via IN and IV routes in only two of the 28 mice PI, but metastasized to other organs (liver, spleen, kidney, small and large intestines) at a higher rate (Figure 7D-E and S8D).

Next, we developed an orthotopic xenograft lung tumor model of HCC4017 from K2 or R5 cells. Mice inoculated orthotopically in the left lung with tumor organoids grown in R5 then pre-treated in K2 HCC4017, died within 60 days of injection (Figure 7F). However, HCC4017 cells constantly maintained in R5 media were sacrificed 180 days post-injection (Figure 7F). Interestingly, HCC4017 K2 and R5 cells both formed tumors with slight but not significant differences in size or number of lung tumors (Figure 7G and S9A). However, we did observe differences in the colonization of organs outside of lung, as well as increased necrosis of primary lung tumors indicating increased capacity for extra-pleural metastases (Figure 7G and S9A).

Lactate production by the cytoplasmic enzyme, lactate dehydrogenase A (LDH) acts as one of the main gate-keepers to altered tumor glycolysis. Not surprising, LDH is differentially expressed in cancer cells compared to healthy tissue, while its expression is hierarchically distributed even among different cancer subtypes from the same organ^60^. We examined four of the tumors formed (from males and females injected with cells grown in R5 and K2 prior). HCC4017 tumors derived from cells grown in high nutrients, regardless of sex, formed heterogenous tumors composed of epithelial and mesenchymal tumor clusters while K2-adapted cells formed tumors with more homogenous morphology like the observations, we initially made in vitro (Figure. 3e). Immunostaining for LDH within these tumors interestingly revealed that the epithelial portions of the tumors derived from the R5 grown cells, had high LDH staining while surrounding mesenchymal/stromal compartments had no LDH expression (Fig. 5e). Cells adapted to K2 formed tumors with uniform expression of LDH (Fig. 5e). From this we were able to conclude our in vivo data correlated with our in vitro data demonstrating low-nutrient conditions promoted increased migration/invasion and metastasis through reprogramming of tumor metabolism toward glycolysis.

Perineural invasion (PNI), observed in many non-neuroendocrine pancreatic adenocarcinomas (80-100%)^61–63^ and recently triple-negative breast cancers^64^, associate PNI with intense pain and high rates of metastasis^61^. In addition to poorer survival, recent studies have also associated PNI with dysregulation of the TME promoting high-lactate metabolism via neural-TME and neural-tumor crosstalk ^62–64^. We observed a 5-fold increase in the presence of PNI per tumor in the low-nutrient derived tumors compared to high-nutrient derived tumors (Extended Fig. 6e). Furthermore, the epithelial compartments of the R5 tumors had high LDH expression, while the PNI did not. K2 tumor-associated PNIs had high expression of LDH and uniform expression of LDH within the tumors (Fig. 5e and Extended Fig. 6e). Taken together these data suggest in aggressive lung tumors where nutrient attainment is low, the occurrence of PNI may be a mode by which tumors are able to increase their glycolytic output as observed in pancreatic tumors but to a lesser extent (Extended Fig. 6e)^63^.

## Discussion

Life is dependent on the fundamental process of cellular locomotion for proper embryonic development, regeneration, and wound healing. While high-throughput screening for compounds targeting cancer cell survival, death mechanisms, and proliferation is successfully awaiting or has been approved by the FDA ^65,66^, compounds that specifically target the metastatic cascade are few and still desperately needed ^67,68^. We initiated our current study to investigate potential therapeutic vulnerabilities in cancer cells from various tissues before, during, and after collective migration. We screened 60+ cell lines derived from normal non-malignant cells, together with primary and metastatic cancer cell lines, in their respective growth media. Our original screen was done to find the best migrators for further investigation. We found that the migration of the non-cancerous cells was significantly greater than the cancer cells regardless of the growth media. Collectively we found there were drastic differences between how the lung cancer cells responded to wounding compared to how the pancreatic and breast cancer cells reacted to wounding. Much of our experimental follow-up focused mostly on malignant/non-malignant lung cells. Non-malignant primary and immortalized epithelial cells from lung, pancreas, and skin tended to act similarly with regards to their: 1) ability to migrate and invade robustly 2) localization of iATP during migration and 3) response to metabolic inhibition. These differences ultimately reflect fascinating tissue-specific biology potentially related to the cells of origin of the individual cell line.

Lung cancer cells adapted to low-nutrient conditions not only increased their migration to same levels as normal cells but also changed their metabolism matching that of the mobile non-malignant cells. Critically, these findings suggest that more attention should be given to the media conditions of in vitro studies, to reveal new targets that are masked by high nutrient conditions. Simultaneously, the heterogenous compositions of both epithelial and mesenchymal cells observed in high-nutrients switched to a homogenous mesenchymal-like composition when grown short-term in low-nutrient conditions. Taken all together, our data suggests during migration, invasion, and most likely metastasis, lung and pancreatic cancer cells utilize different avenues of glucose metabolism, whereby lung cells had a propensity to metabolize glucose to lactate, while the pancreatic cancer cells seemingly carried glucose through the TCA/OxPhos cycles (Figure 5g). Whether this is a phenomenon dependent on the cells of origin and oncogenotype remains to be seen.

Cancer has been described as “wounds that do not heal” based on the observations of H.F. Dvorak in 1986, comparing the similarities of fibroblast-derived ECM proteins that are secreted during normal wound healing and tumor formation^69^. During normal homeostatic wound repair, discrete epithelial stem cell populations contribute to wound repair via balancing proliferation, differentiation, and migration at distinct regions within the wound^8^. Epithelial cells need calcium to survive and fully differentiate in order to form and maintain cell–cell adhesion junctions^70^. However, a fine balance is required as primary non-malignant lung cells grown under the same conditions as cancer cells undergo senescence, abnormal proliferation, and/or survival constraints^49,71,72^. During normal keratinocyte wound repair (as well as with other epithelial cell types) concentrations of calcium have been shown to physiologically lower in response to damage and during the repair process ^73^. Even so, we found short-term R5 acclimated normal lung epithelial cells (with double the calcium levels) still closed wounds faster than lung cancer counterparts. Thus, we suggest conventional cancer growth media conditions are not suitable for in vitro experiments investigating migration/invasion. We anticipate the fallout of our findings will be two-fold: 1) A paradigm shift away from cell migration assays conducted in supraphysiological-nutrient growth media and 2) a shift toward better in vitro modeling of in vivo metastatic events.

## Methods

### Cell lines and treatment

Lung cancer cells (unless otherwise noted) and human bronchial epithelial cells (HBECs), un-immortalized and immortalized, were derived and maintained by the Minna lab in Hamon Cancer Center, UTSW Dallas, TX^49,71,72^. The immortalized human pancreatic ductal epithelial (HPDE H6c7) cell line was purchased from Kerafast (EAC001-FP). Human pancreatic cancer cell line PANC1 and human pancreatic adenocarcinoma cell line CAPAN-1 were obtained from the Brekken lab (UTSW, Dallas, TX). C5LM2 was a cell line derived from liver metastasis from a PANC1 tumor-bearing mouse and generated by the Brekken lab^74^. Human breast cancer cells, HCC1954 Parental and the, LCC2 – isolated from brain-metastases– were obtained from the Malladi lab (UTSW, Dallas, TX)^75^. Human triple-negative breast cancer (TNBC) cells MDA-MB-231, human ER-positive breast cancer cells MCF-7, and human mammary epithelial cell line MCF10A were obtained from the Martinez lab in the Hamon Cancer Center, UTSW Dallas, TX. Human keratinocyte cell lines HaCaT and hTERT-expressing human keratinocytes were obtained from the Harris-Tryon lab (UTSW Dallas, TX). Small cell lung cancer patient-derived cells, MGH1505-1A3, was obtained from Drapkin lab (Hamon Cancer Center, UTSW, Dallas, TX). Human embryonic stem cell line H9 was prepared by Dr. Sushama Sivakumar (UTSW Dallas, TX). Human colorectal cancer (CRC) cell lines HCT116, DLD1, RKO, LOVO and HT-29 and immortalized human colonic epithelial cell HCEC1CT were provided by Whitehurst lab (UTSW, Dallas, TX). Immortalized human colonic epithelial cell HCEC1CT was established by Shay lab^76^ and maintained by the Whitehurst Lab (UTSW, Dallas, TX). Human melanoma cell lines SK-MEL-2, SK-MEL-37, A375, WM3211, UACC257 and MNT1 and immortalized melanocytes and human prostate cancer cell lines PSK1, ARCaP, C4-2B, DU145, PC3, LNCaP and 22Rv-1 and human prostate epithelial cell line RWPE-1 were maintained by the Whitehurst lab (UTSW Dallas, TX). All lung, breast, pancreas, skin cancer cell lines and A375 were cultured in RPMI Sigma Aldrich, R8758) with 11mM glucose and 2mM glutamine supplemented 5% FBS (26140-079, Gibco / 35-015-CV, Corning), SK-MEL-37, UACC257, LNCaP, 22Rv-1, ARCaP and C4-2B were cultured in RPMI, Sigma Aldrich, R8758) with 11mM glucose and 2mM glutamine supplemented 10% FBS (26140-079, Gibco / 35-015-CV, Corning) and all colorectal cancer cell lines, MNT1, SK-MEL-2, WM3211, DU145 and PC3 were grown in advanced DMEM, 12491-015, Gibco supplemented 10% FBS (26140-079, Gibco / 35-015-CV, Corning) and maintained at 37 °C in an atmosphere of 5% CO_2_. Human bronchial epithelial cell lines, HPDE H6c7, HK-2, RWPE-1 and hTERT Keratinocytes were maintained in keratinocyte-SFM (KSFM) media supplemented with Bovine Pituitary Extract (BPE) and human recombinant Epidermal Growth Factor (rEGF) per manufacturer’s instruction (Cat. No. 17005042, Thermo Fisher Scientific). MCF10A was cultured in MEGM^TM^ Mammary Cell Growth media per manufacturer’s instruction (CC-3150, Lonza). Human melanocytes were maintained in Medium 254 with HMGS. HCEC1CT culture method was as described^76^, without modification. Human cell lines were authenticated to confirm origin, and all cell lines were pre-treated with Plasmocin (ant-mpt-1, InvivoGen) and all cell lines were confirmed to be free of mycoplasma (Myco-Sniff-Rapid^TM^ Mycoplasma Luciferase Detection Kits, 3050402, MP Biomedicals) before use.

### Adaptation of cancer cells to Keratinocyte-SFM

Cancer cells in Figure 1 were all adapted for 8-10 passages until they repeatedly were able to exponentially growth with > 90% cell viability in RPMI with 11 mM glucose and 2 mM glutamine supplemented with 5% FBS. Further short-term (3 days) and long-term (2 weeks) adaptation of cancer and normal cells were done in the human plasma-like medium (HPLM, cat. no. A48991-01 LOT no. 2502529)^40^ alone, in 2% or 5% FBS, and KSFM alone or with 2% FBS.

### High-throughput scratch wound cell migration assay with imaging

For all scratch wound cell migration assays, 50,000 cells were seeded onto bovine collagen (125-50, CELL APPLICATINS INC) or Matrigel-coated 96-well microplates and incubated at 37 °C with 5% CO_2_ incubator for 24 hours. Homogeneous linear scratches were made using IncuCyte^®^ 96-well Wound-maker (4563, Sartorius). The wounded cells were washed 3 times with PBS pH 7.4 to remove the detached cells and then 100 μL of medium was added. Images of the wounds were automatically acquired within the CO_2_ incubator by IncuCyte^®^ SX5 live-cell analysis system (Sartorius). Images were captured every 1-2 hours for the duration of the experiment. Images were analyzed by defining a scratching mask and a cell confluence mask using a software algorithm on the IncuCyte^®^ Scratch Wound Analysis Software Module (9600-0012, Sartorius).

### High-throughput scratch wound invasion assay with imaging

For wound invasion an open wound area was created in the cell monolayer using the IncuCyte^®^ 96-well Wound-maker (4563, Sartorius) and washed 3 times with PBS pH 7.4 to remove the detached cells. After removing any detached cells, use a pre-chilled cool box to equilibrate the plate at 4°C for 5 minutes before carefully aspirating the cold medium. Diluted Matrigel^®^ Growth Factor Reduced (GFR) was added to the appropriate wells and placed into the cell culture incubator for 30 minutes. Then 100 μL of medium was added to the appropriate wells and place the plate into the IncuCyte^®^ SX5 live-cell analysis system (Sartorius). Images were captured every 2 hours for the duration of the experiment. Images were processed by defining cell density of the wound using a software algorithm. Data analysis was performed using the IncuCyte^®^ Scratch Wound Analysis Software Module (9600-0012, Sartorius).

### Label-free real-time cell proliferation assay

Cells were seeded at 2×10^3^ cells per well in 96-well, clear-bottomed, tissue culture plates in 100 μL complete growth medium and left to equilibrate at room temperature for 30 minutes before 37 °C, 5% CO_2_ incubator. After overnight incubation, fresh KSFM was replaced and the plate was inserted into the IncuCyte SX5 live-cell analysis system (Sartorius) for real-time imaging, with four fields imaged per well under 10x magnification every 2 hours for a total of 72 hours. Data were analyzed by defining a cell confluence mask using a software algorithm.

### Immunofluorescence staining

About ∼50,000 cells were grown on 35mm glass-bottom cover dishes (P35G-1.5-14-C, MatTek) first coated with bovine collagen solution (125-50, CELL APPLICATINS INC) for 1 hour. Wound inductions at T=0 were made using pipette-suction method. Cells were rinsed in PBS twice and fixed with 4% paraformaldehyde diluted in PBS pH 7.4 for 1 hour at room temperature. Cells were permeabilized with 0.5% Triton X-100 in PBS for 5 minutes and incubated in blocking solution (5% Donkey serum and 0.5% Triton X-100 in PBS) for 30 minutes at room temperature. Cells were incubated with the following primary antibodies at concentrations of 1:500 or 1:250 diluted in blocking solution: Vimentin (Mouse mAb, ab8978-Abcam, or Goat pAb, PA5142829-Invitrogen), Tubulin (Rabbit pAb, ab18251-Abcam). The following day, cells were washed with PBS 3 times and then incubated with secondary antibodies in blocking solution for 30 minutes from Invitrogen (Donkey anti-mouse Alexa Flour 488, Donkey anti-Goat IgG2a Alexa Flour 488, Donkey anti-Rabbit Alexa Fluor 568, Phalloidin 568 and DAPI). Cells were washed with PBS 3-5 times before imaging.

### Western blot analysis

Cells were harvested after PBS washes in RIPA Lysis Buffer (89901, Thermo Fisher Scientific) supplemented with protease inhibitor cocktail (78425, Thermo Fisher Scientific), followed by sonication and centrifugation. Protein concentration was measured by the bicinchoninic acid (BCA) kit (23225, Thermo Fisher Scientific), resolved on 4-20% Mini-PROTEIN^â^TGX^TM^ Precast Protein Gels (4561093, BIO-RAD), and transferred to a 0.2mm nitrocellulose membrane. The membrane was blocked for 30 minutes at room temperature in 1x TBS with 0.1% Tween 20 (TBS-T) containing 5% BSA, then probed with primary antibodies. Immunoblotting was performed with primary antibodies against human, Anti-**β** Actin (CST / #3700) Anti-Hexokinase I (CST / #2024), Anti-Hexokinase II (CST / #2867), Anti-GAPDH (CST / #5174), Anti-a-Tubulin (CST / #3873S), Anti-LDHA (CST / #3582), Anti-Pyruvate Dehydrogenase (CST / #3205), Anti-ATP5B (SCBT/ # sc-55597) Image acquisition was performed using the LI-COR Odyssey Fc.

### Intracellular ATP measurement using IncuCyte

For measurement of intracellular ATP level in live cells using IncuCyte ATP-binding FRET indicator, cell lines stably expressing a genetically encoded, fluorescent ATP sensor or a control (non-ATP-binding) sensor were generated by IncuCyte CytoATP Orange Lentivirus Reagent Kit (4772, Sartorius). Cell ATP levels were monitored over days and analyzed using the IncuCyte SX5 Live-Cell Analysis System configured with an SX5 Metabolism Optical Module (4820, Sartorius). Conformational changes take place upon ATP binding by employing a dual excitation (535X, mKok orange; 485X, cpmEGFP green). For the quantitative assessment, the IncuCyte ATP Analysis Software Module (9600-0033, Sartorius) allowed ATP-dependent changes to calculate the image’s ratio of fluorescence intensity (485X | 578M) / (535X | 578M), enabling direct measurement of ATP independent of cell number or indicator expression level.

### High-throughput screen of real-time ATP levels during short-term adaptation and migration

Intracellular ATP stable cell lines were seeded in bovine collagen or Matrigel-coated 96-well microplates at the density of 30,000 cells per well and incubated overnight at 37 °C in 5% CO_2_ incubator. At sub-confluence, cancer cells were adapted to 2% FBS were or continuously grown in RPMI with 11 mM glucose and 2 mM glutamine supplemented with 5% FBS while normal cells were continuously grown in KSFM with 2%FBS or adapted to RPMI with 11 mM glucose and 2 mM glutamine supplemented with 5% FBS for 3 days (short-term adaptation). Wounds were made in confluent monolayer via mechanical tool, IncuCyte 96-well Wound-maker (4563, Sartorius), During migration, high-resolution images were captured subsequently to monitor cell morphology, real-time ATP levels and the dynamics of cell migration using the IncuCyte SX5 for Metabolism specialized optical module and image acquisition mode. Images were analyzed to measure ATP demands in the IncuCyte ATP Analysis Software Module.

### Nutrient utilization rates

To measure metabolic rates, cells were plated into 6-well plates. At confluent density, an open wound area was created in the cell monolayer using the pipette-suction method, rinsed in PBS once and fed with 5mL of RPMI with dialyzed 5% FBS, KSFM or KSFM with 2% FBS. Conditioned medium was collected at 0, 4 and 24 hours after wound induction. Concentrations of glucose, lactate, glutamine, and glutamate were determined from 1mL aliquots of medium using an automated electrochemical analyzer (BioProfile Basic-4 analyzer; NOVA). Metabolic rates were determined by normalizing absolute changes in metabolite abundances to final protein content.

### Seahorse XF analysis

All oxygen consumption and extracellular acidification measurements were conducted using an Agilent Seahorse XF24 Analyzer (Agilent Technologies). HCC4017 and PANC1 parental cells were plated in 24-well XF Cell Culture Microplate (100777-004, Agilent Technologies). Next day, the medium was removed and cells were rinsed in 1C PBS. RPMI with 5% FBS or KSFM with 2% FBS was added to respective wells and cells were incubated for 3 days (Short-term adaptation). On the day of the assay, culture media was removed and replaced with Seahorse assay media. Cells were allowed to acclimate to assay media for 1 hour in the incubator, then wounds were induced using the pipette-suction method. Once acclimated, the cells were assayed for basal OCR, ATP-dependent OCR, and maximal OCR using a standard mito-stress kit protocol. For the assay, we used 10uM oligomycin and 2uM CCCP for the injections. Once the assay was complete, cells were collected. Total protein levels were measured by the bicinchoninic acid (BCA) kit (23225, Pierce Biotechnology) to normalize ensure the observed changes in respiration do not manifest from changes in cell number.

### Metabolomics screening and analysis

#### Extraction of Metabolome from cell lines

Cells were changed the medium of the cell plates 2 hours before metabolite extraction. The cells were rinsed with ice-cold normal saline solution once and then aspirated. Cells were taken at each time point and immediately placed on dry ice (−80°C). 500μL of 80% (vol/vol) acetonitrile in water, pre-cooled in −80°C freezer for at least 1 hour. The dishes on dry ice were transferred to the −20°C and left for 5 minutes. Cells were scraped into the extraction solvent on dry ice (−80°C). The solution was transferred to Eppendorf tubes, and frozen in liquid nitrogen. The solution was subjected to three freeze-thaw cycles between liquid nitrogen and 37°C. After the third thaw, the solution was vortexed for 1 minute and then centrifuged at 20000xg for 15 minutes in a refrigerated centrifuge. The metabolite-containing supernatant was transferred to a new tube and kept the protein pellet to perform protein quantification.

Metabolite abundance data were acquired from a targeted metabolomics screening and imported into R (v4.x) for all downstream analyses. The dataset comprised 336 metabolites measured across three cell lines: HBEC30UI (Un-immortalized human bronchial epithelial cell), HCC4017 in R5, and HCC4017 adapted to K2. For each cell line, metabolite abundances were quantified under a control condition (Ctrl) and at two post-treatment time points (T=0 and T=4), with two biological replicates per condition.

#### Data preprocessing and quality control

The raw abundance matrix contained 0.28% missing values and 16.51% zero entries. Zero values were treated as below the limit of detection and recorded as missing (NA). Metabolites for which all replicates within any single cell line were missing (i.e., more than four NA values per cell line group) were removed, reducing the dataset from 336 to 257 metabolites, with 4.67% residual missingness.

#### Normalization and imputation

The filtered data were first normalized by total ion current, whereby each sample’s metabolite abundances were divided by the corresponding column sum to correct for differences in total signal intensity across samples. The remaining missing values were then imputed using the missForest algorithm, a nonparametric, random-forest–based method applied to the transposed data matrix. Following imputation, variance-stabilizing normalization (VSN) was applied using the normalized VSN function from the vsn/limma Bioconductor packages to stabilize the variance across the intensity range.

### Animal Experiments (Xenograft models)

All animal-related experiments and procedures were performed with the Guide for Care and Use of Laboratory Animals approved by the Institutional Animal Care and Use Committee (IACUC) at UTSW. Athymic nude mice (6902, Inotiv Envigo) from 8 to 24 weeks of age were used for studies, housed in pathogen-free conditions under the supporting care of the Animal Resource Center (ARC).

### Part 1 – SubQ

To generate tumors, HCC4017 in RPMI with 5% FBS (R5) or adapted for 3 days to KSFM with 2% FBS (K2) medium were injected subcutaneously at 500k, or 1×10^6^ cells/mouse. To generate pancreatic cancer tumors, PANC1 PA spheroids in RPMI with 5% FBS (R5) or pre-conditioned short-term to KSFM with 2% FBS (K2) medium was injected subcutaneously at 1×10^6^ cells/mouse. Subcutaneous injections were done by anesthetizing the mouse with isoflurane via open drop and with a 27 ½ gauge needle, 50 μL of cell suspension was injected into the right flank. Mice were monitored for recovery as well as for tumor formation and measured using a digital caliper (62379-531, Control Company) once a week for quantification.

### Part 2 – Lung Orthotopic Injections

To generate lung cancer tumors, previous lung tumor organoids generated from SubQ HCC4017 in RPMI with 5% FBS (R5) or short-term adapted to KSFM with 2% FBS (K2) medium diluted in 25% ECM gel (growth factor reduced)/PBS solution was injected orthotopically at 500k cells/mouse. Lung orthotopic injections were done following the Principles of Aseptic Technique and Rodent Survival Surgery per the ARC at UT Southwestern and IACUC policies #128.5 (Analgesia) and #218.9 (Anesthesia). Sections 3.3 Orthotopic Injection and 3.4 Removal of Wound Clips of a previous paper (Alexandra N. Wilson et al. 2022) describing this procedure were followed with a few modifications. Syringes (U-100 Insulin Syringe, REF 26014, Comfort Point) and cell mixtures were kept on ice in a sterile hood.

The mouse was transferred from the anesthesia chamber into a sterile hood placing the mouse in the right lateral recumbent position and kept under anesthesia. Buprenorphine Extended Release at 0.5-1.0mg/kg was administered subcutaneously before prepping the mouse for surgery. Once the mouse was prepped, confirmation that the mouse was anesthetized was done by checking the toe pinch reflex before continuing. The syringe was loaded with cells using a P200 pipette. The cell mixture was resuspended, and 25 μL was loaded onto the syringe by removing the plunger and dispensing the cell mixture into the open end of the syringe. The plunger was then reinserted, making sure that any air bubbles were removed by tapping the syringe and moving the plunger back and forth. The prepped syringe was then placed nearby on a sterile Kim wipe. Following that, the lower edge of the mouse ribs and the scapula were located and, using forceps, the skin of the left lateral thorax was pulled up and away from the chest wall. Using surgical scissors, an approximately 1 cm transverse incision approximately 0.5 cm below the inferior border of the left scapula along the midline long axis was made. Additional incisions through the subcutaneous fat and fascia were made until the chest wall was visualized taking care of avoiding nicking any visible blood vessels. The respiring lung was identified, a pale pink structure under the rib cage. To inject the needle just above rib 6, the lowermost rib, which should be rib 8, was identified. The ribs were visually counted cranially until ribs 6 and 5 were reached. Using a 70% ethanol wipe to sterilize the gloved index and middle fingers, the incision area was held open applying light pressure to the chest wall, and the needle was inserted 4 mm into the midline of the lateral side at a 90-degree angle to the chest wall. Then, the cells were slowly injected into the lung holding the needle in place for a few seconds after injection to ensure successful cell transplantation. It’s important to note that lung parenchyma may be visible immediately after injection. The incision site was closed by lining up the underlying tissue and skin and placing 2 to 4 wound clips with an Autoclip applicator (Autoclip Wound Closing System, Cat # NC9946451, Fisher Scientific). The mouse was then placed alone in a warmed recovery cage and monitored until fully recovered from anesthesia before returning to a cage with other mice. Additional monitoring for recovery was done every 12 hours for the next 72 hours following the surgery. After 10 – 14 days, the wound clips were removed by anesthetizing the mouse and removing the wound lips with an Autoclip remover. Tumors were traced by noninvasive bioluminescence (BLI) imaging of experimental animals using AMI-HTX (Spectral Instruments Imaging) as well as weekly weigh-ins.

Citation for protocol followed: Wilson AN, Chen B, Liu X, Kurie JM, Kim J. A Method for Orthotopic Transplantation of Lung Cancer in Mice. Methods Mol Biol. 2022; 2374:231-242. doi: 10.1007/978-1-0716-1701-4_20. PMID: 34562257; PMCID: PMC9262117.

### Part 3

#### Animal Experiments (Xenograft models) Part 1 – IV / IN

To generate tumors, luciferase-labeled HCC4017 at 2e10^6 cells/mouse in RPMI with 5% FBS (R5) or pre-conditioned for three days to KSFM with 2% FBS (K2) medium were injected intravenously via tail vein (I.V.) and intranasally (IN). HCC1954 breast cancer cells (250k cells/mouse) were injected via the tail intravenously. Intravenous injections via the tail were done by using a 27 ½ gauge needle, at a max volume of 0.125ml for 25g mouse, and into the lateral tail vein with the assistance of a tail illuminator (supplied by the ARC). Intranasal instillation of cancer cells was performed by anesthetizing the mouse with isoflurane via open drop. Using a 200P pipette, 25 μL of cell suspension was instilled into each nostril dropwise (50 μL total volume). Mice were monitored for recovery, and tumors were traced by noninvasive bioluminescence (BLI) imaging of experimental animals using AMI-HTX (Spectral Instruments Imaging). Bioluminescence images were quantified using the ROI tool in Aura software. Subcutaneous tumors were monitored and measured once a week for quantification.

### Tumor organoids culture

The following media formulations were used in the protocol: 3D culture media was prepared from Advanced DMEM (12491015, Gibco)/Ham’s F-12K (30-2004, ATCC) medium supplemented with 1′B-27 supplement (17504044, Gibco), 1′Insulin-Transferrin-Selenium (ITS-G) supplement (41400045, Gibco), Penicillin-Streptomycin (30-002-CI, Corning), 1′GlutaMax Supplement (35050061, Gibco) and 1mM HEPES (H3375, Sigma Aldrich); Digestion media was prepared from Advanced DMEM/F-12K medium with 20% FBS and contained 5mg/mL Collagenase/Dispase (10269638001, Roche) and 100 mg/mL DNase I (10104159001, Roche). Stop solution against TrypLE was prepared from Advanced DMEM/F-12K with 20% FBS.

### Generation of tumor organoids

Tumor tissue was placed in a 15mL conical tube containing 10mL PBS pH 7.4 (10010-023, Gibco) with 20mg/mL gentamicin (15710-072, Gibco) (PBS-G) on ice for transport. Tumors were washed three times with cold PBS-G and minced into 1-2mm diameter pieces in digestion buffer with a sterile scalpel: digestion buffer contains 5mg/mL Collagenase/Dispase (10269638001, Roche) and 100 mg/mL DNase I (10104159001, Roche). Digestion mix was transferred into a 50mL falcon tube and incubated at 37 °C in 5% CO_2_ incubator for 1 hour. Every 15 minute during the incubation period, the suspension was vortexed gently. The supernatant was discarded, and 3D culture media was added to the fragment to wash, then centrifuged at 1200 rpm for 5 minutes to pellet. The pellet was incubated with TrypLE (12604021, Gibco) at 37 °C in 5% CO_2_ incubator for 10 minutes and neutralized with 20% FBS in AdDMEM/F-12K. Mixture was passed through a 100-mm strainer (22363549, Fisher Scientific) to remove large undigested tissue clusters. It was centrifuged at 1200 rpm for 5 minutes to remove supernatants, and pellet was washed with 3D culture media. The cell pellet was resuspended and counted to collect 1.0 ′ 10^5^ for being embedded in 300 mL of Matrigel (354230, Corning; Growth Factor Reduced). Matrigel-cell mixture was dispensed as 50-mL droplets into one well of a 6-well plate. The drops were solidified for 1 minute right side up and for another 9 minutes upside down at 37 °C in 5% CO_2_ incubator. After the drops solidified, 3mL of warm 3D culture media was added to each well.

### Histopathology and immunohistochemistry

Formalin-fixed tissue samples were sent to the Tissue Management Shared Resource to obtain Hematoxylin and Eosin (H&E) stained sections as well as paraffin-embedded (FFPE) tumor sections of 4-mm in thickness. Specific protein expressions were visualized by immunohistochemistry (IHC) for further analysis. Briefly, tissue sections were de-paraffinized with xylene (5 minutes, twice) then rehydrated through an ethanol/PBT (PBS with 0.1% (vol/vol) Tween-20) gradient (3-minute washes in each of 100% (twice), 95%, 70%, 50%, and PBT). The hydrated tissue sections were boiled for 30 minutes at 106°C - 110°C using a TintoRetriever pressure cooker (BSB7087, BioSB) in diluted ImmunoDNA Retriever 20X with citrate solution (BSB0022, BioSB), (Final 5 ′ concentration). After the tissue sections cooled down, they were washed in PBT for 5 minutes before blocking for 20 minutes with blocking serum (PBT: Donkey Serum (D9663-10ML, MilliporeSigma), 10% Triton X-100, Glycine 1M (1610718, BIO-RAD)). The tissue sections were incubated with primary antibodies and DAPI (DAPI dilution (1:10,000)-Stock DAPI concentration 1 mg/mL) diluted in blocking serum at 4°C overnight. Sections were mounted with Aqua-Poly/Mount (18606-20, Polysciences) and analyzed by microscope (BZ-X810, Keyence).

### Statistics and reproducibility

Each exact n value and the number of independent experiments is indicated in each figure legend. Standard deviations of the mean were added to the appropriate graphs and all statistical tests mentioned in this manuscript were performed by GraphPad Prism (Version 10.4.2) software. Immunofluorescence staining data were from three independent experiments to ensure reproducibility, and similar results were obtained in Supplemental Figure 5. Tests of reproducibility were performed by several lab members across weeks and months for the migration/invasion screens in low-and high-nutrient conditions, and mouse models of colonization/metastasis. All malignant and non-malignant cells were thawed monthly with routine testing for mycoplasma performed.

## Acknowledgements

We acknowledge the Kim Laboratory for training with the orthotopic lung experiments, all present and past members of the Osborne Lab and current members of the Cobb and Whitehurst Lab and Mike Peyton for cells and reagents. We also acknowledge Joe Albanesi, Rolf Brekken, and our Beloved David Mangelsdorf for critical feedback on all versions of manuscript. We thank Dionne Ware for administrative help. We acknowledge the assistance of the University of Texas Southwestern Tissue Management Shared Resource, a shared resource at the Simmons Comprehensive Cancer Center, which is supported in part by the National Cancer Institute under award number P30 CA142543. We acknowledge the assistance of the Children’s Medical Center Research Institute (CRI), and Data Science Shared Resource (DSSR) at UT Southwestern Medical Center for metabolomics screening and analysis, respectively. Initial funding for this project included CPRIT (Cancer Prevention Research Institute of Texas) start-up grant number, RR210016 and the V Foundation grant number V2021-028. Current funding: NIH/NIGMS 1R35GM160161. S.F.L. was supported by Cancer Prevention and Research Institute of Texas (CPRIT) Pre-doctoral Training Grant number, RP210041. B.K. and I.T. were partially funded by CPRIT Post-doctoral Training Grant number, RP210041.

