## Supplemental Figures for "Media matters: A resource for understanding the determinants of migratory potential of malignant and non-malignant cells"

#### Supplental Figure 1. Non-malignant Cells Close Their Wounds Faster Than Malignant Cells

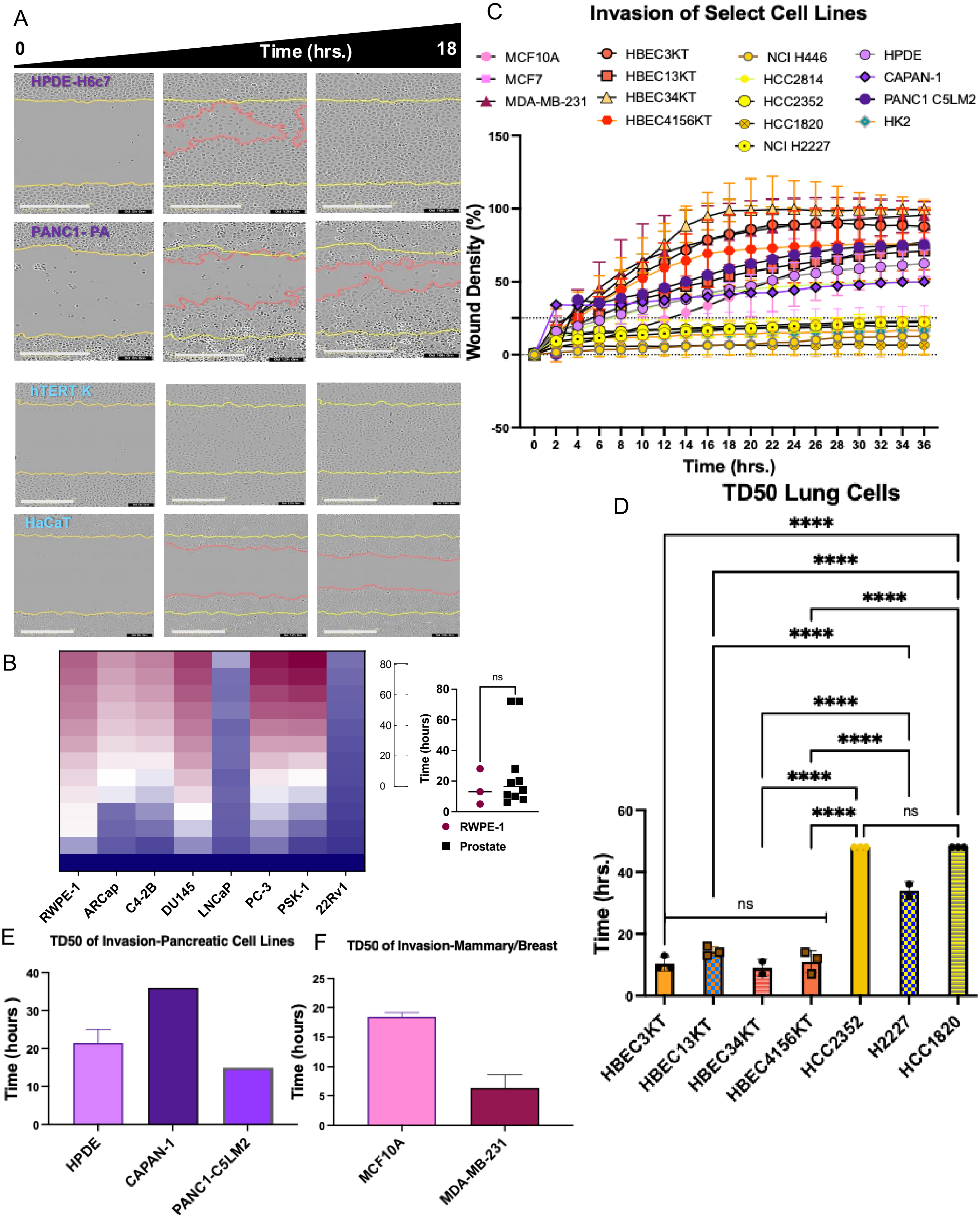

**Supplemental Figure 1. Non-malignant Cells Close Their Wounds Faster Than Malignant Cells.** A) Representative images of normal (HPDE H6c7, hTERT Keratinocyte; hTERT K) and cancer cells (PANC1-PA, HaCaT) from pancreas and skin. Yellow lines in images indicate the initial wounded area; red lines represent cells moving into the wound. Scale bars, 400mm. B) Heatmap (Left) shows the migratory rate of normal and cancer cells and dot plot graph (Right) represents the quantification of the time for wound of normal and cancer cells to reach at 80% in prostate. Statistical significance assessed using unpaired t-test, benign prostate epithelial cell line RWPE-1, N=3 and all prostate cancer cell lines, N=10. ns=0.5002. C) Scratch wound invasion graph of wound density in seven non-malignant cell lines; MCF10A (breast), HBEC3KT, HBEC13KT, HBEC34KT, HBEC4156KT (lung), HPDE H6c7 (pancreas), and HK (kidney) and nine malignant cell lines; MCF7, MDA-MB-231 (breast), NCI H446, HCC2814, HCC2352, HCC1820, NCI H2227 (lung), CAPAN-1, PANC1 C5LM2 (pancreas). Error bars are from individual experiments, N=2. Dotted line represents cut-off of cells ability to invade (<25%). Graphs measuring TD50 to assess cellular invasiveness of non-malignant and malignant cells from D) lung, E) pancreas F) breast. All error bars indicate the mean ± SD of N=3 independent experiments for d) and N=2 independent experiments for e) and f). * Statistically significance of lung cell lines *(P < 0.05) ** (P < 0.005) ***(P < 0.0005) **** (P < 0.0001), ns means not significant. Statistical significance was assessed using one-way ANOVA for panel D).

**Supplemental Table 1. Table of cell lines tested for their migratory and invasive capacity**

| Cell Lines Tested for Migration/Invasion | | | | | |
|---|---|---|---|---|---|
| Normal (non-tumorigenic) cell lines | | | Cancer (tumorigenic) cell lines | | |
| **Cell Line** | **Migrate** | **Invade** | **Cell Line** | **Migrate** | **Invade** |
| HBEC3KT | Yes | Yes | A549 | Yes | No |
| HBEC13UI | Yes | Yes | HCC1313 | Yes | N/A |
| HBEC13KT[Δ] | Yes | Yes | NCIH2227* | Very Slow | No |
| HBEC14UI | Yes | N/A | NCIH446 | Yes | Slow° |
| HBEC14KT | Yes | N/A | NCIH460[Δ] | Slow | No |
| HBEC30UI[a] | Yes | Yes | NCIH520[Δ] | Slow | No |
| HBEC30KT[a] | Yes | Yes | HCC1820[Δ] | Slow | Slow |
| HBEC34KT[b] | Yes | Yes | HCC2352[AA] | Yes | No |
| HBEC56KT[AA] | Yes | Yes | HCC2814 | Yes | Yes |
| HBEC4156UI[AA] | Yes | Yes | HCC4017[a] | Yes | Yes |
| HBEC4156KT[AA] | Yes | Yes | HCC4018[b] | No | No |
| MCF10A | Yes | Yes | NCIH1993[c] | Slow | N/A |
| HPDE-H6c7 | Yes | Yes | NCIH2073[c] | Yes | N/A |
| HK-2 (Kidney) | Slow | Yes° | NCIH2085[d] | Yes | Yes |
| hTERT Keratinocyte | Yes | N/A | NCIH2086[d] | Yes | Slow° |
| HCEC1CT | Yes | Yes | MGH1505-1A3 | Yes | N/A |
| Melanocytes (immortalized) | Yes | No | MDA-MB-231 | Yes | Yes |
| RWPE-1 | Yes | Yes | MCF7 | Yes | Yes |
| | | | HCC1954-Parental[Δe SA] | Yes | Yes |
| | | | HCC1954- LCC2[e SA] | Yes | Yes |
| | | | CAPAN-1 | Yes | Yes |
| | | | PANC1-Parental[f] | Yes | Yes |
| | | | PANC1-C5-LM2[f] | Yes | Yes |
| | | | HaCaT | Yes | Yes |
| H9 (hESC) | No | No | HCT116 | Yes | Yes |
| | | | DLD1 | Yes | Yes |
| | | | RKO[AA] | Yes | Yes |
| | | | LOVO | Yes | Yes |
| | | | HT-29 | Yes | Yes° |
| | | | SK-MEL-2 | Yes | Yes |
| | | | SK-MEL-37 | Yes | Yes |
| | | | A375 | Yes | Yes |
| | | | WM3211 | Yes | Yes |
| | | | UACC257 | Yes | Yes |
| | | | MNT1 | Yes | Yes |
| | | | PSK1 | Yes | Yes |
| | | | ARCaP | Yes | Yes |
| | | | C4-2B | Yes | Yes |
| | | | DU145 | Yes | Yes |
| | | | PC3 | Yes | Yes |
| | | | LNCaP | Yes | Yes |
| | | | 22Rv-1 | Yes | Yes |

KT- CDK4/hTERT overexpression; UI un-immortalized/primary

[a, b, c, d, e, f] Isogeneic Cell lines from same patient

[AA] Black/African decent, [SA] South Asian decent

°- Did not invade/migrate well

**Lung;Breast/Mammary;Skin/Keratinocyte;Pancreas;Colon; Melanoma; Prostate**

N/A= not tested for invasion

Δ= change in morphology during migration

'*'= Adherent cell line, originally characterized as mixed adherent and suspension

**Supplemental Table 1. Table of cell lines for their migratory and invasive capacity.**

Table of non-tumorigenic/normal and tumorigenic cell lines screened for the ability to migrate on collagen and invade Matrigel. H9-human embryonic stem cells were plated to confluence on hESC Matrigel (Corning 354277). Cancer cell lines were derived from primary and metastatic biopsied tumors of lung, pancreas, breast, skin, colon, melanoma/melanocytes and prostate including three isogeneic pairs. The lung isogeneic pair, NCIH2085/H2086, was isolated from a primary and metastatic lymph node tumor from the same cancer patient. The other two pairs, pancreas and breast, were initiated orthotopically in immune-compromised mice using human tumor cell lines (PANC1 parental and HCC1954 parental) then allowed to metastasize (PANC1-C5LM2 isolated from a liver metastases / HCC1954-LCC2 isolated from a latent brain metastases). Non-malignant/normal cells were also isolated from the same tissues mentioned above. Normal lung biopsies were first established as primary un-immortalized cultures that were eventually immortalized via overexpression of hTERT and CDK4 (HBEC3KT, HBEC13UI/13KT, HBEC14UI/KT, HBEC30UI/KT, HBEC34KT, HBEC56KT, HBEC4156UI/KT). Other isogeneic series were isolated from a biopsied lung tumor HCC4017, normal adjacent lung tissue was used to generate the non-malignant un-immortalized HBEC30UI, then the immortalized HBEC30KT cell lines. Five human colon cancer cell lines and one human colonic epithelial cell line were included in the present study. HCEC1CT is an immortalized human colonic epithelial cell (HCEC) lines with hTERT and CDK4. Colorectal cancer cells were HCT116, DLD1, RKO, LOVO and HT-29. Six melanoma cell lines SK-MEL-2, SK-MEL037, A375, WM3211, UACC257 and MNT1 and normal human immortalized melanomacytes were tested their migratory and invasive capacity. All the prostate cancer (PSK1, ARCaP, C4-2B, DU145, PC3, LNCaP and 22Rv-1) and benign prostate epithelial cell line, RWPE-1 were screened for their migratory and invasive ability. N/A=not tested for Invasion, [D]=change in morphology during migration KT- CDK4/hTERT overexpression; UI un-immortalized/primary [a, b, c, d, e, f,] Isogeneic Cell lines from same patient [SA] South Asian Decent [AA] Black/African decent ; ° Migrated and/or Invaded very slowly *NCIH2227 was originally characterized as a mixed suspension and adherent cell line but, was subcloned to select for adherent cells.

**Supplemental Figure 2. ATP is differentially localized in cancer and non-cancerous cells**

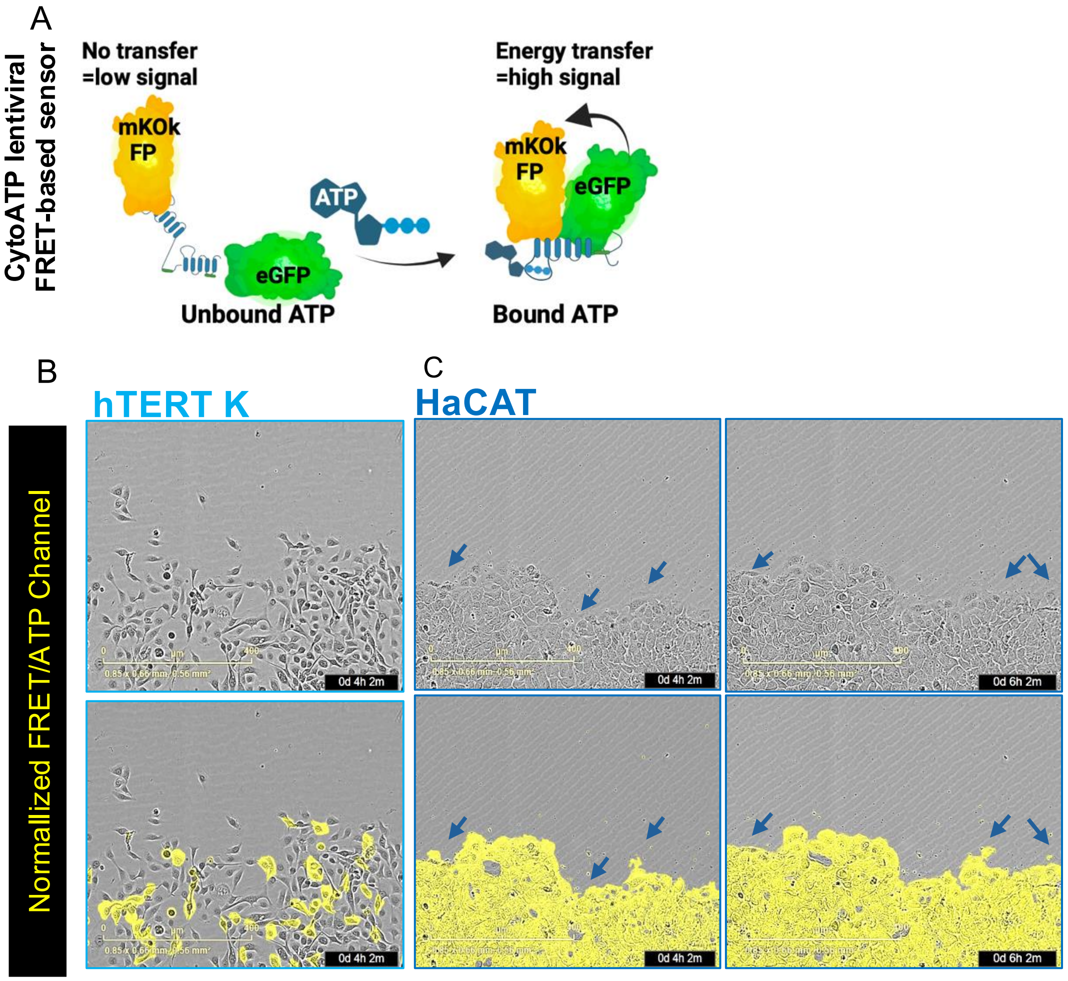


A) Illustration of how intracellular ATP (iATP) dynamics are detected by the FRET-based CytoATP sensor used on the Incucyte®. Wounds were induced in all wells using Incucyte® 96-well wound maker then analyzed with the SX5 metabolism module and data analysis software. Time-lapse visualization of the cells, B) hTERT K and and C) HaCAT during early time points of migration showing dynamic iATP-localized to the membrane ruffles of cancer but not normal cells. Arrows (black and blue) mark iATP localization to the membrane ruffles of cancer cells. Representative images from duplicate wells of N=4 independent experiments. Panel A) made in BioRender.

**Supplemental Figure 3. Quantification of migration rates in various media and serum concentrations**

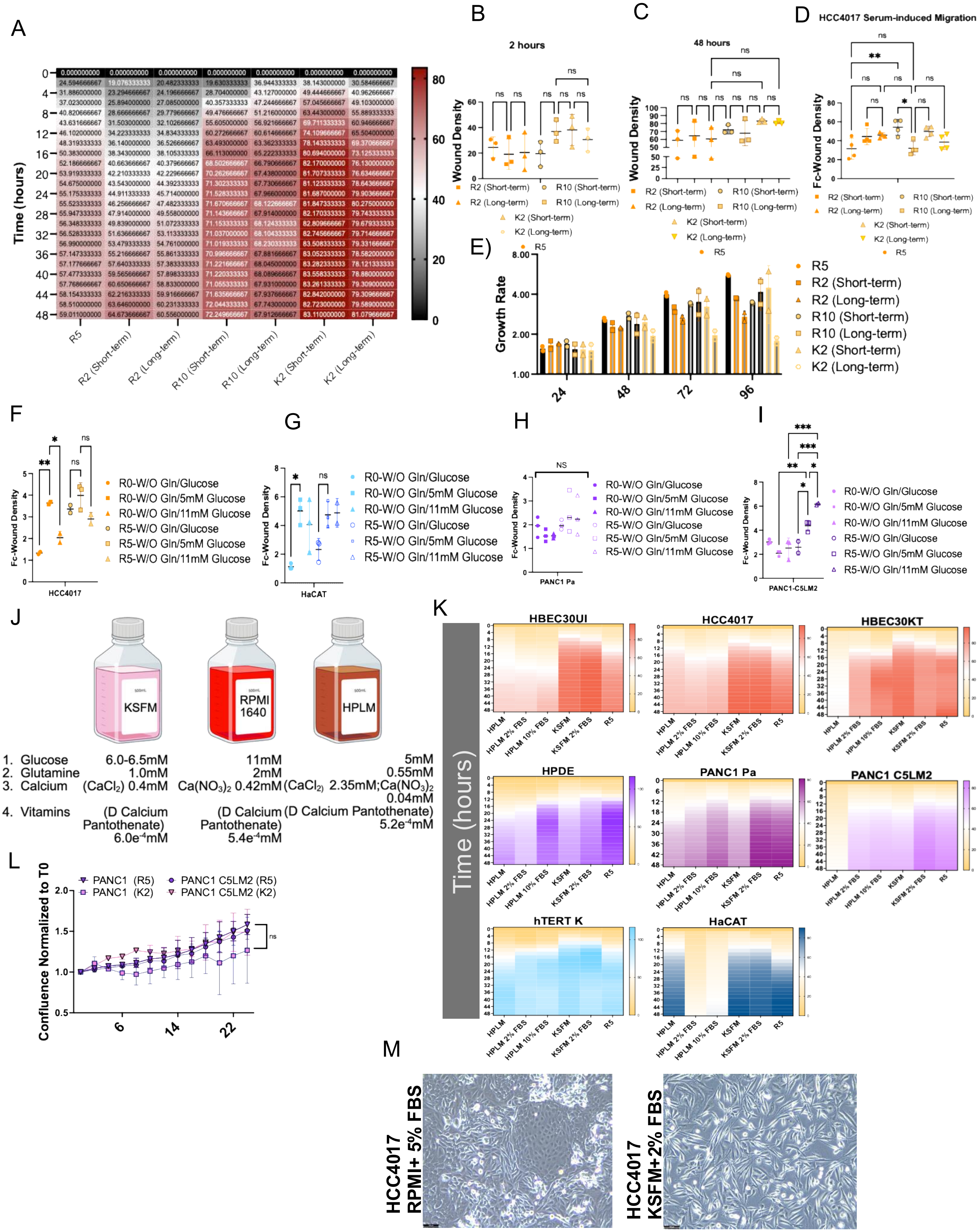

**Supplemental Figure 3. Quantification of migration rates in various media and serum concentrations**

A) Heatmaps showing the wound density of cells adapted short- and long-term to RPMI160 2%, 10% FBS and KSFM 2% FBS. Graphs showing the quantification of wound density of B) at 2 hours C) at 48 hours D) fold-change (Fc) from 2-72 hours. Statistical significance for B) and C) achieved by one-way ANOVA of, N=3 individual experiments. Statistical significance for D) achieved by one-way ANOVA of N=4 individual experiments(**=0.0092, *=.0123). E) Graphs showing the growth rate of N=2 individual experiments.
F-I) Quantification of cells adapted short-term to RPMI1640 (R0) or (R5) without glutamine or glucose then treated with 5mM or 11mM glucose F) HCC4017 (lung) G) HaCAT (skin) H) PANC1 (Pancreas) I) PANC1-C5LM2 (Pancreas). Statistical significance for F-I) achieved by one-way ANOVA of N=3 individual experiments HCC4017 ***<0.001, **<0.005, *=.0275; HaCAT *<0.04; PANC1 n.s >0.300; PANC1-C5LM2 *<0.02, **<0.009, ***<0.0003, ****<0.0001 J) Schematic of various medias tested and deviations from the components of the basal keratinocyte-serum free media (KSFM; not including the additional EGF and bovine pituitary extract). Panel made in BioRender. K) Heatmaps showing the wound density of cells adapted short-term to human plasma-like media + 2% and 10% FBS (HPLM), KSFM +/- 2%FBS (K2 ) or maintained in RPMI 1640 5%FBS (R5). L) Graphs showing the growth rate of pancreatic cancer cells PANC1 and PANC1-C5LM2 adapted short-term in K2 or maintained in R5 media relative to time zero. Statistical significance achieved by two-way ANOVA of individual experiments, N=2 for Panc1 PA and C5LM2 (ns=0.998 for PA; ns=0.8413 for C5LM2). M) Representative images to show different cell morphology by phase contrast of lung malignant HCC4017 in RPMI1640 with 5% FBS and HCC4017 long-term adapted to KSFM with 2% FBS.

#### Supplemental Figure 4. Invasion of lung cells by scratch-wound invasion

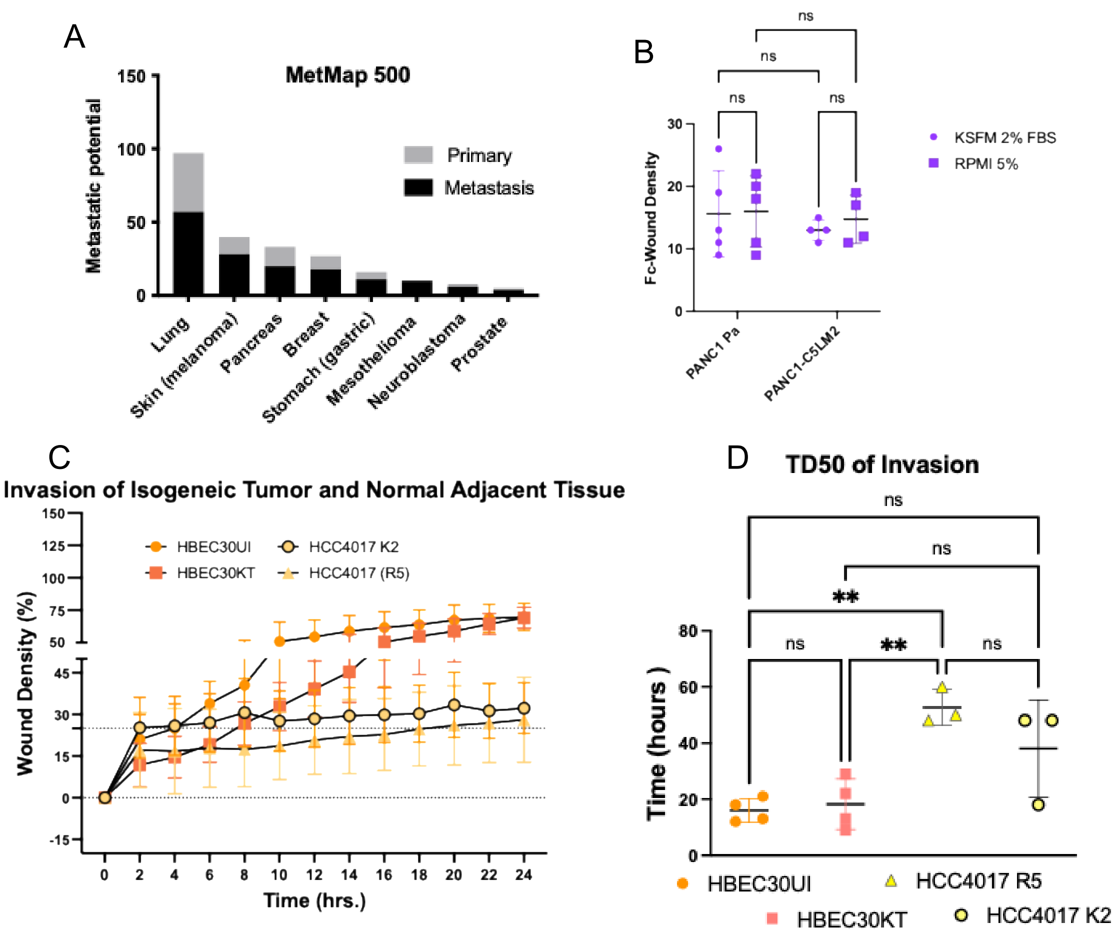


**Invasion of lung cells by scratch-wound invasion**

Movies Of Cancer and Normal Cells invading Matrigel

A) Data exploration of subsets of MetMap500 cell lines sequenced post-intracardial inoculation described in manuscript by Jin, X et al. 2020. B) Graphs showing the quantification of fold-change (Fc) in wound density from 2-24 hours. C) Wound density graph to compare the ability of cells to move through Matrigel in the isogenic lung cell lines HCC4017, HBEC30UI and HBEC30KT from the same patient. HCC4017 K2 was conditioned long-term (two-weeks) in K2. Error bars are from individual experiments (N=3) and represent SD. D) TD50 Quantification of panel C). Error bars are from individual experiments, N=4 for HBEC30UI and HBEC30KT. Statistical significance using one-way ANOVA is from individual experiments. **P < 0.00549 and ns>0.341.

**Supplemental Figure 5. iATP changes in response nutrient changes.**

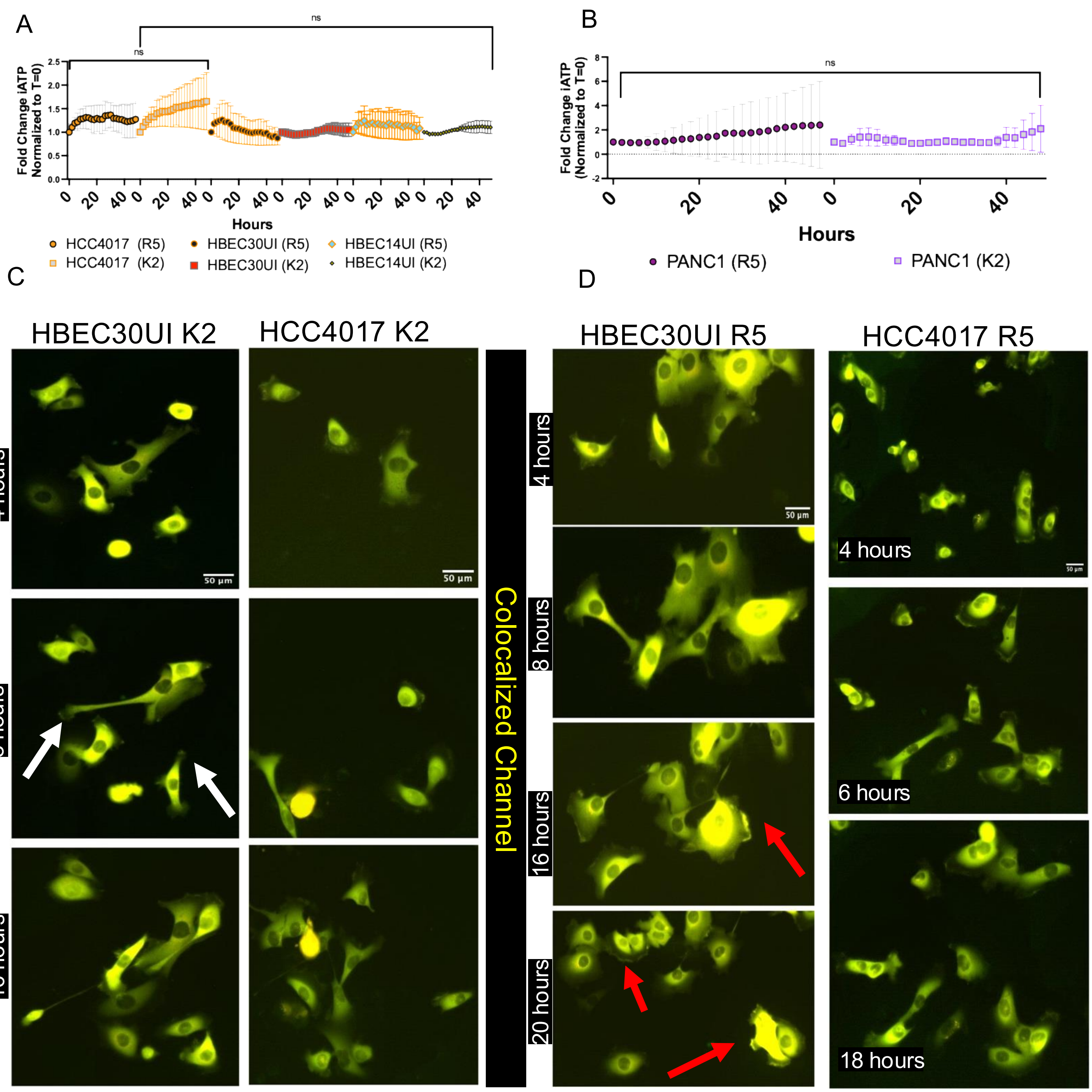


**Supplemental Figure 5. iATP changes in response nutrient changes.**
Fold change in iATP of A) lung cells and B) pancreatic cells measured in Figure 4G) and 4H) for migration. Statistical significance was assessed using two-way ANOVA P=ns C) and D) Images of cyto-ATP expressing HBEC30 and HCC4017 cells plated sub-confluently to images membrane dynamics of the iATP construct grown in K2 and R5.

**Supplemental Figure 6. Hierarchical clustering of metabolites during in vitro migration**

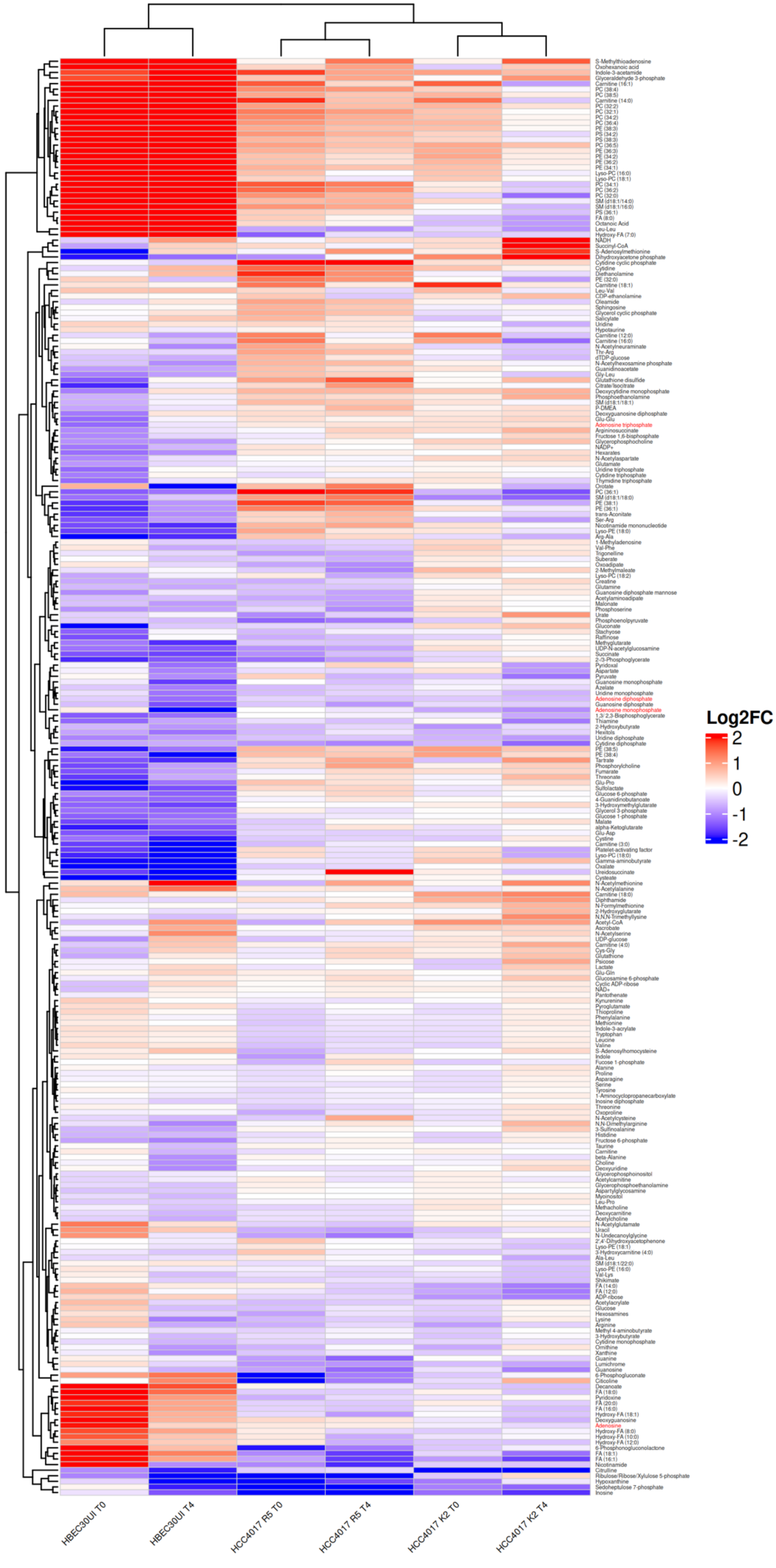

### Supplemental Figure 7. Expression analysis of genes required for glucose metabolism

A

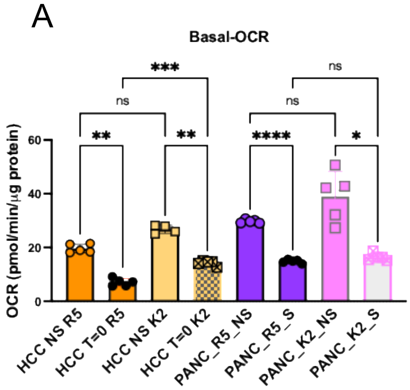


B

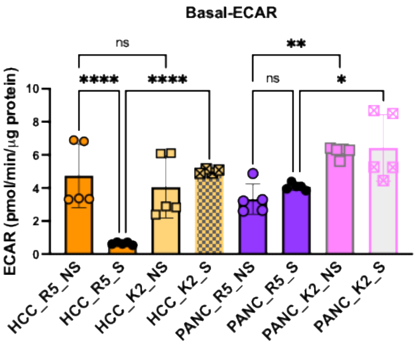


C

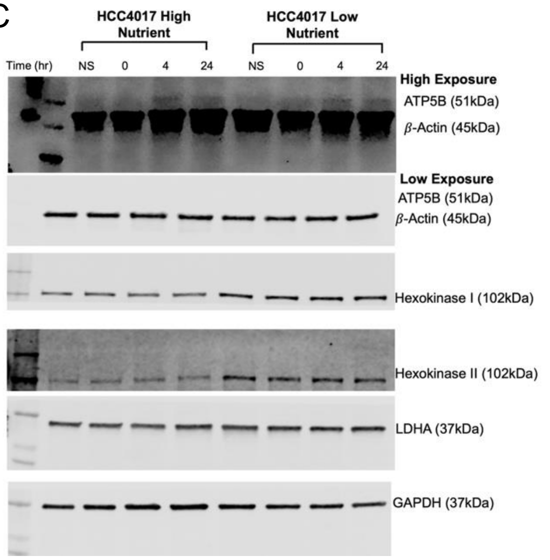


D

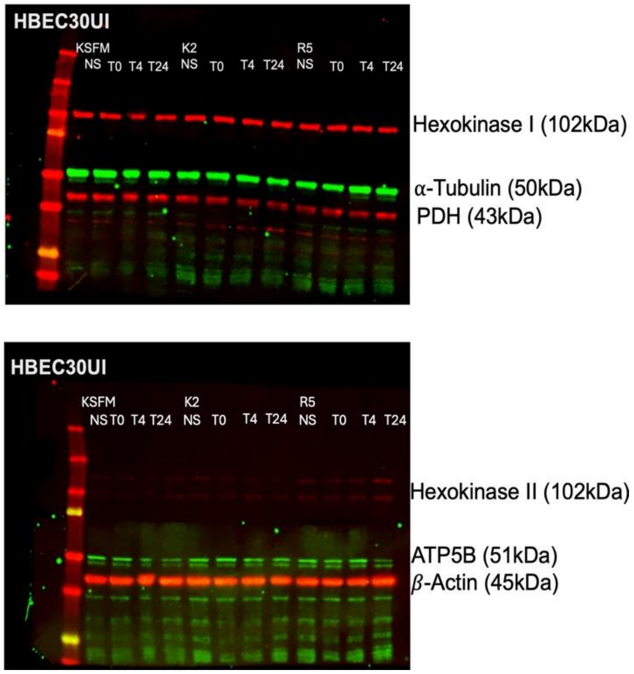


E

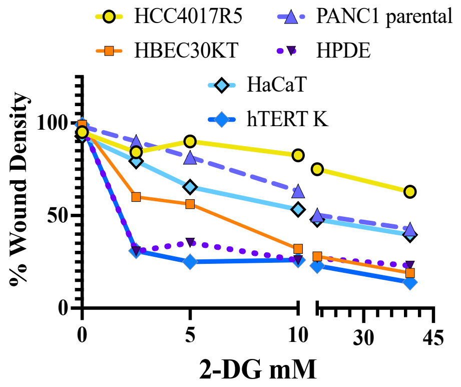


F

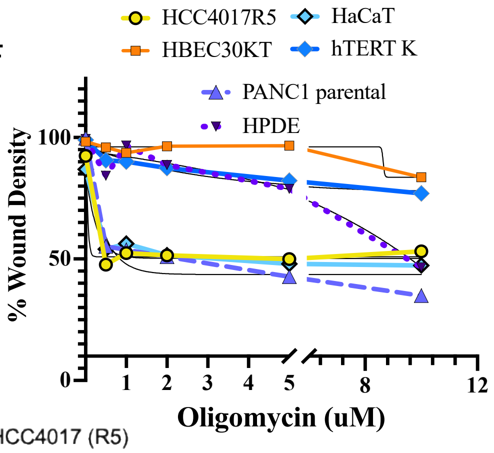


G

| Cell lines | 2DG IC50 (mM) | Oligomycin IC50 (uM) |
|---|---|---|
| HCC4017 | 58.38 | 2.3 |
| HBEC30KT | 5.5 | 58.62 |
| PANC1 PA | 22.05 | 1.83 |
| HPDE | 2.3 | 12 |
| HaCaT | 14.58 | 2.4 |
| hTERT K | 1.79 | 26.11 |

H

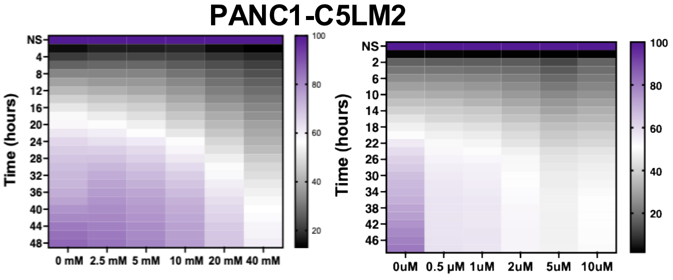


I

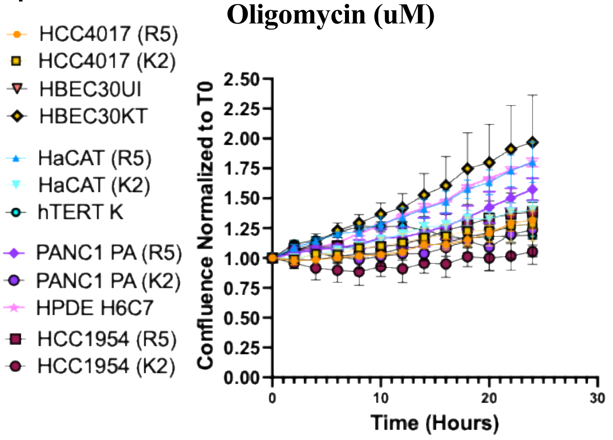

**Supplemental Figure S7. Expression analysis of genes required for glucose metabolism**

Relative A) Basal-OCR and B) Basal-ECAR normalized to total protein before and after wounding of PANC1 and HCC4017 cells in R5 or K2 media. Statistical significance for A) and B) achieved by one-way ANOVA of individual experiments, N=4 or N=5 individual experiments. Basal OCR – P*= 0.0495, **<0.006, *** = 0.0009, ****<0.0001; Basal ECAR –P*<0.048, **<0.0064, ****<0.0001. C) Immunoblots showing levels of ATP5B, Hexokinase I, Hexokinase II, LDHA and GAPDH from HCC4017 in high nutrient and low nutrient environment during the migratory phase (NS: Non-Scratch, T=0-, 4- and 24-hours post-wound induction). b-Actin used as an endogenous control. Representative of N=3 individual experiments. D) Representative western blot to compare protein expression of Hexokinase I, Hexokinase II, ATP5B and PDH from lung non-malignant HBEC30UI in different nutrient environments during the migratory phase (NS: Non-Scratch, T=0-, 4- and 24-hours post-wound induction). b-Actin and a-Tubulin have been used as an endogenous control for protein expression. Representative of N=3 individual experiments. E) IC50 of 2DG and F) oligomycin for the ability to fill in the wound– measuring TD50 (time for wound density to reach 50%). G) Table of IC50 values 2DG and oligomycin in normal and cancer H) Heatmaps quantifying the % wound density post induction over the course of 48hrs of treatment of wounded PANC1-C5LM2 metastatic cells treated with 2DG (left panel) and oligomycin (right panel) (2-Deoxy-D-glucose 2.5 mM, 5 mM, 10 mM, 20mM, 40mM, and Oligomycin 0.5M, 1mM, 2mM, 5mM, 10mM). N=4 individual experiments. I) Graphs showing the growth rate of lung (HCC4017, HBEC30), skin (hTERT K, HaCAT), Pancreas (PANC1 PA, HPDE), and breast (HCC1954) in R5 and K2.

**Supplement To Figure 7. Figure S8. Cells Grown In Low Nutrients Form More Metastases**

A)

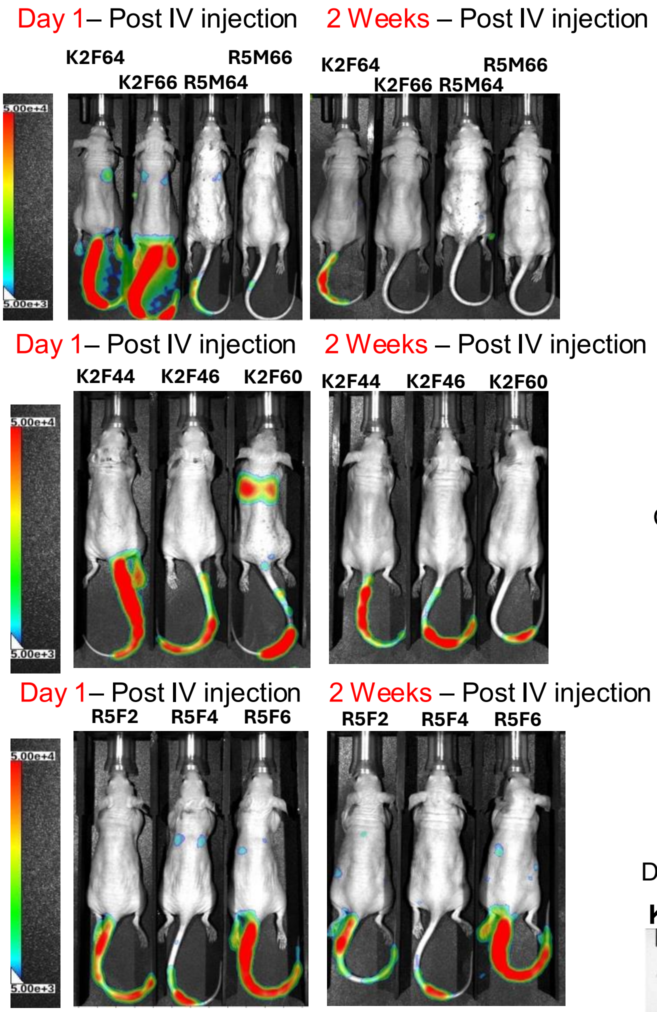


B)

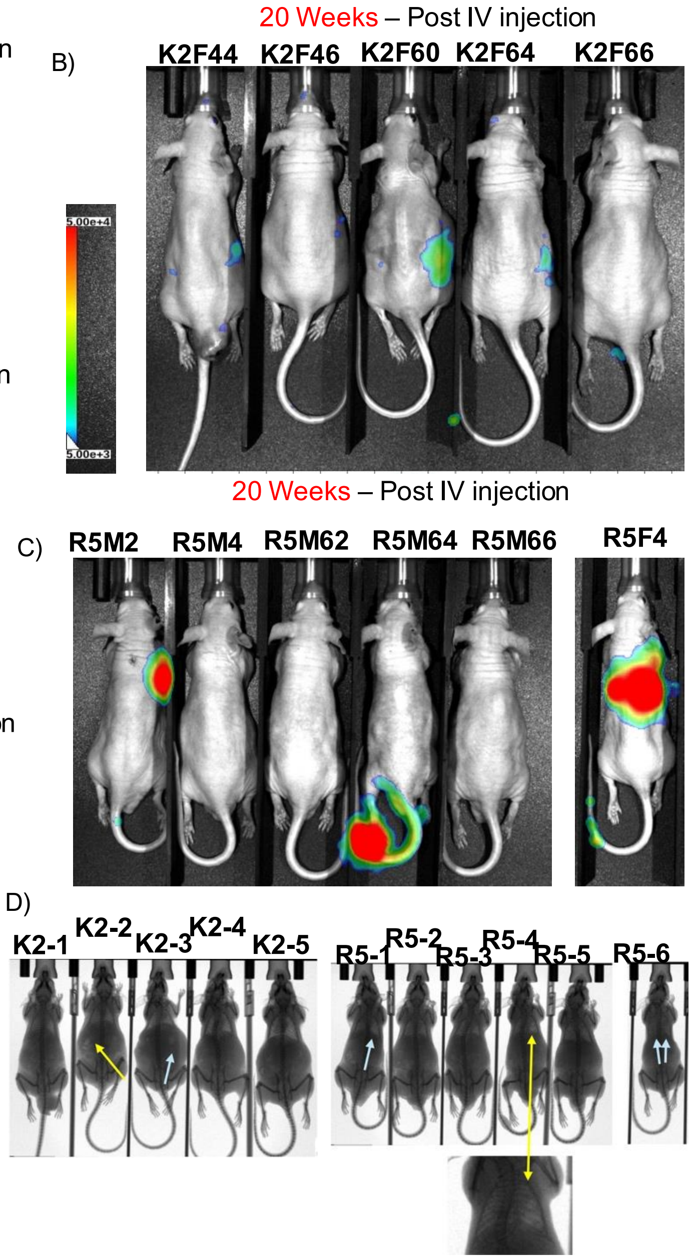


E)

**Tumor Formation (IV, SubQ, IN)**

| | Total mice used | Mice # w/tumors | Rate of formation (%) |
|---|---|---|---|
| R5 | 33 | 7 | 21% |
| K2 | 34 | 11 | 32% |

**No. of mice with tumors via IV-tail vein or intranasal instillation**

| | HCC4017-R5 | | HCC4017-K2 | |
|---|---|---|---|---|
| **Total tumors** | 4 out of 24 | 16% | 7 out of 25 | 28% |
| **Metastases** | 1 out of 24 | 4% | 4 out of 25 | 16% |

**Supplemental Figure 8. Cells Grown In Low Nutrients Form More Metastases**
Cells were infected transiently with an adenoviral luciferase construct. 2e6 HCC4017 cells administered by IV tail vein from adherent cultures in Nude mice. n=16 mice–R5, N=8; K2, N=8. A-C) Mice were imaged on day 1 , day 14, and 20 weeks post-injection (PI) for luciferase. D) At 20 weeks PI mice were also x-rayed for visualization of lung tumors. E) Tables showing tumor formation and number of metastases present from IV model.

**Supplement to Figure 7 Figure S9 . Highly glycolytic lung tumors have increased mortality rates in mice and humans**

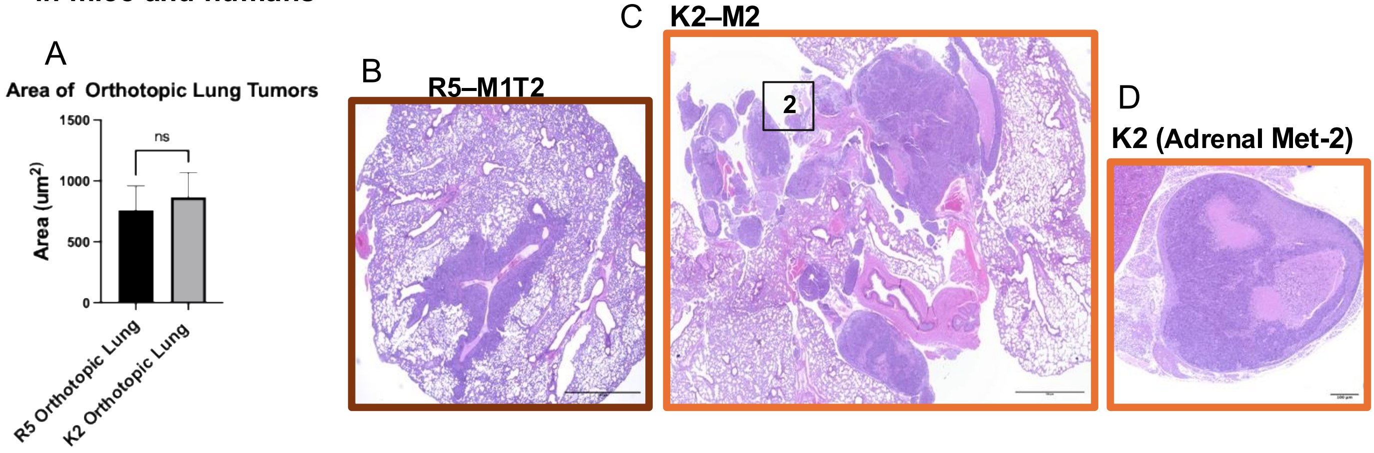


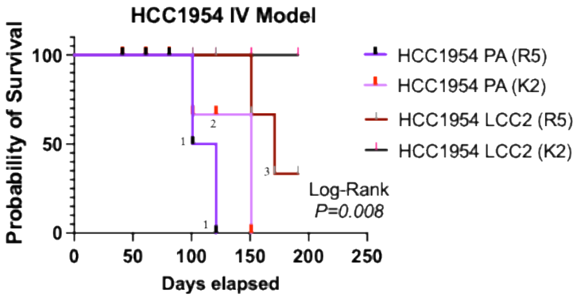


1) Epithelial neoplasms colonized in all lobes of lung
2) Epithelial neoplasms colonized all lobes of lung and necrotic lesions found in brain
3) Epithelial neoplasms colonized all lobes of lung

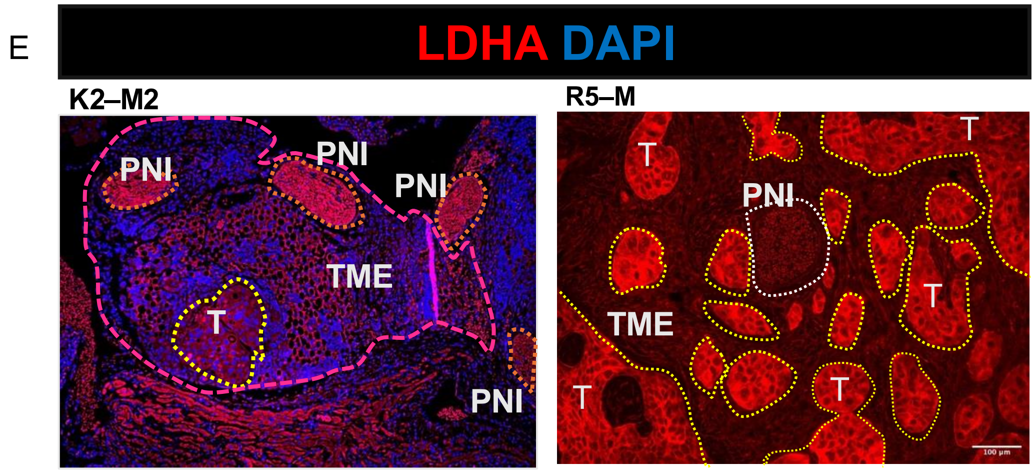


**Supplemental Figure 9 . Highly glycolytic lung tumors have increased mortality rates in mice and humans** a) Quantification of tumors sizes from two independent orthotopic lung models in Figure 7G using Welch's t test, ns=0.789. B-D) Tumor-burdened lungs of a males from K2-adapted and R5 from were formalin-fixed, sectioned and stained via IHC and IHC-IF and for B-D) H&E and E) lactate dehydrogenase (LDH) and DAPI. LDH staining in tumors with epithelial morphology (demarcated with yellow dotted lines), tumor microenvironment and perineural invasion (PNI, demarcated with orange dotted lines in K2-derived mouse and white dotted lines in R5-derived mouse
